# Studies of DNA ‘breathing’ by polarization-sweep single-molecule fluorescence microscopy of exciton-coupled (iCy3)_2_ dimer-labeled DNA fork constructs

**DOI:** 10.1101/2023.09.27.559819

**Authors:** Jack Maurer, Claire S. Albrecht, Patrick Herbert, Dylan Heussman, Anabel Chang, Peter H. von Hippel, Andrew H. Marcus

## Abstract

Local fluctuations of the sugar-phosphate backbones and bases of DNA (often called DNA ‘breathing’) play a variety of critical roles in controlling the functional interactions of the DNA genome with the protein complexes that regulate it. Here we present a single-molecule fluorescence method that we have used to measure and characterize such conformational fluctuations at and near biologically important positions in model DNA replication fork constructs labeled with exciton-coupled cyanine [(iCy3)_2_] dimer probes. Previous work has shown that the constructs that we test here exhibit a broad range of spectral properties at the ensemble level, and these differences can be structurally and dynamically interpreted using our present methodology at the single-molecule level. The (iCy3)_2_ dimer has one symmetric (+) and one anti-symmetric (–) exciton with respective transition dipole moments oriented perpendicular to one another. We excite single molecule samples using a continuous-wave linearly polarized laser with polarization direction continuously rotated at the frequency 1 MHz. The ensuing fluorescence signal is modulated as the laser polarization alternately excites the symmetric and the anti-symmetric excitons of the (iCy3)_2_ dimer probe. Phase-sensitive detection of the modulated signal provides information about the distribution of local conformations and conformational interconversion dynamics of the (iCy3)_2_ probe. We find that at most construct positions that we examined the (iCy3)_2_ dimer-labeled DNA fork constructs can adopt four topologically distinct conformational macrostates. These results suggest that in addition to observing DNA breathing at and near ss-dsDNA junctions, our new methodology should be useful to determine which of these pre-existing macrostates are recognized by, bind to, and are stabilized by various genome regulatory proteins.

## 1. Introduction

The assembly of the protein-DNA complexes that drive the various processes of genome expression involves coordinated interactions between multiple protein-nucleic acid complexes. For example, during the initial stages of DNA replication, proteins recognize and bind to base-sequence-specific sites at the replicative origin, forming a multi-subunit protein-DNA complex that, in turn, recruits additional protein factors to establish the ‘trombone’ shaped framework of the functional DNA replication-elongation complex [1]. This DNA framework is subject to thermally induced fluctuations that permit local regions of the initially double-stranded (ds) DNA to transiently expose single-stranded (ss) DNA templates to the aqueous surround, and thus provide access to the protein components of the DNA replication complex. Although the notion of ‘DNA breathing’ is consistent with free energy landscape models of protein-DNA interactions [2], little is known about the distributions of the Boltzmann-weighted conformational macrostates at and near replication fork junctions, or the associated activation barriers that control the assembly and operation of replication protein-DNA complexes.

In this paper we describe a novel, polarization-selective single-molecule fluorescence method to study the local conformations and conformational dynamics of model DNA fork constructs that are labeled with pairs of cyanine dyes [(iCy3)_2_], which have been rigidly inserted into the sugar-phosphate backbones at several defined positions at and near the ss-dsDNA fork junction (see Fig. 1*A* and 1*B*) [3]. The closely spaced iCy3 monomers (labeled *A* and *B*) are electrostatically coupled, so that the (iCy3)_2_ dimer supports delocalized excitons [symmetric (+) and anti-symmetric (−)] with orthogonally polarized electric dipole transition moments (EDTMs),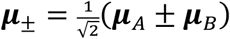. The magnitudes of the EDTMs depend sensitively on the local conformation of the (iCy3)_2_ dimer probe, which is characterized by the ‘tilt’ angle, θ_*AB*_, the ‘twist’ angle, *ϕ*_*AB*_, together with the distance between the two iCy3 monomers, *R*_*AB*_ (see Fig. 1*C*) [4-7].

**Figure 1.**
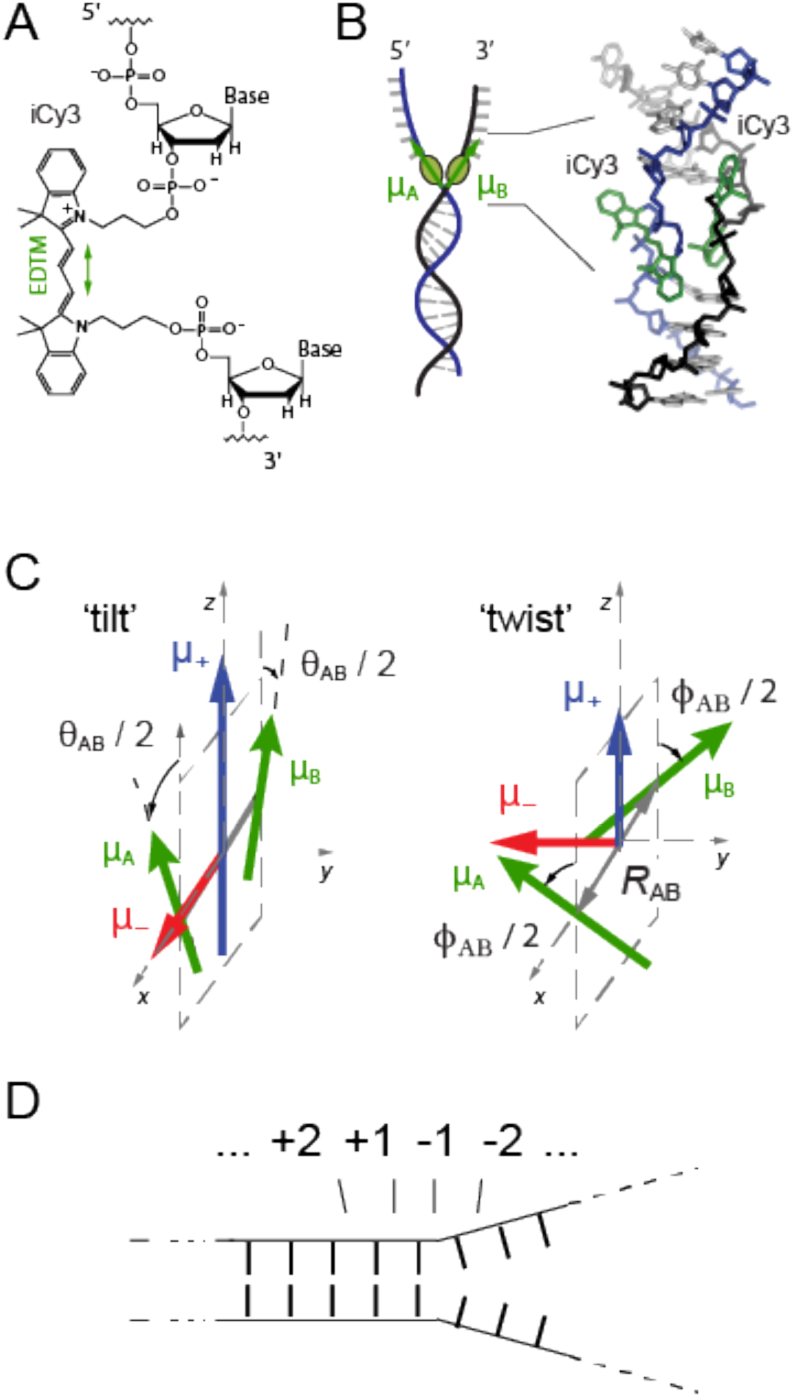
Labeling chemistry and nomenclature of the internal (iCy3)_2_ dimer probes positioned within the sugar-phosphate backbones of model ss-dsDNA constructs. (***A***) The Lewis structure of the iCy3 chromophore is shown with its 3’- and 5’-linkages to the sugar-phosphate backbone of a local segment of ssDNA. The double-headed green arrow indicates the orientation of the electric dipole transition moment (EDTM). (***B***) An (iCy3)_2_ dimer-labeled DNA construct contains the dimer probe near the single-stranded (ss) – double-stranded (ds) DNA junction. The conformation of the (iCy3)_2_ dimer probe reflects the local secondary structure of the sugar-phosphate backbones at the probe insertion site position. The sugar-phosphate backbones of the conjugate DNA strands are shown in black and blue, the bases in gray, and the iCy3 chromophores in green. (***C***) The structural parameters that define the local conformation of the (iCy3)_2_ dimer probe are the inter-chromophore separation vector *R*_*AB*_, the tilt angle θ_*AB*_, and the twist angle *ϕ*_*AB*_. The electrostatic coupling between the iCy3 chromophores gives rise to the anti-symmetric (−) and symmetric (+) excitons, which are indicated by the red and blue arrows, respectively, and whose magnitudes and transition energies depend on the structural parameters. (***D***) The insertion site position of the (iCy3)_2_ dimer probe is indicated relative to the pseudo-fork junction using positive integers in the direction towards the double-stranded region, and negative integers in the direction towards the single-stranded region.

In Fig. 2, we show a schematic layout of our experimental approach, which we call polarization-sweep single-molecule fluorescence (PS-SMF) microscopy. A single (iCy3)_2_ dimer-labeled DNA construct is resonantly excited using a continuous-wave (cw) laser whose plane polarization is rotated at the frequency Ω/2π = 1 MHz. The symmetric (+) and anti-symmetric (−) excitons of the (iCy3)_2_ dimer probe, which are oriented perpendicular to one another, are alternately excited as the laser polarization direction is rotated across the corresponding EDTMs, and the ensuing modulated fluorescence is detected using the phase-tagged photon-counting (PTPC) method [8]. The resulting PS-SMF signal is directly related to the local conformation of the (iCy3)_2_ dimer-labeled probe within the ss-dsDNA construct, the fluctuations of which can be monitored on time scales of microseconds or longer.

**Figure 2.**
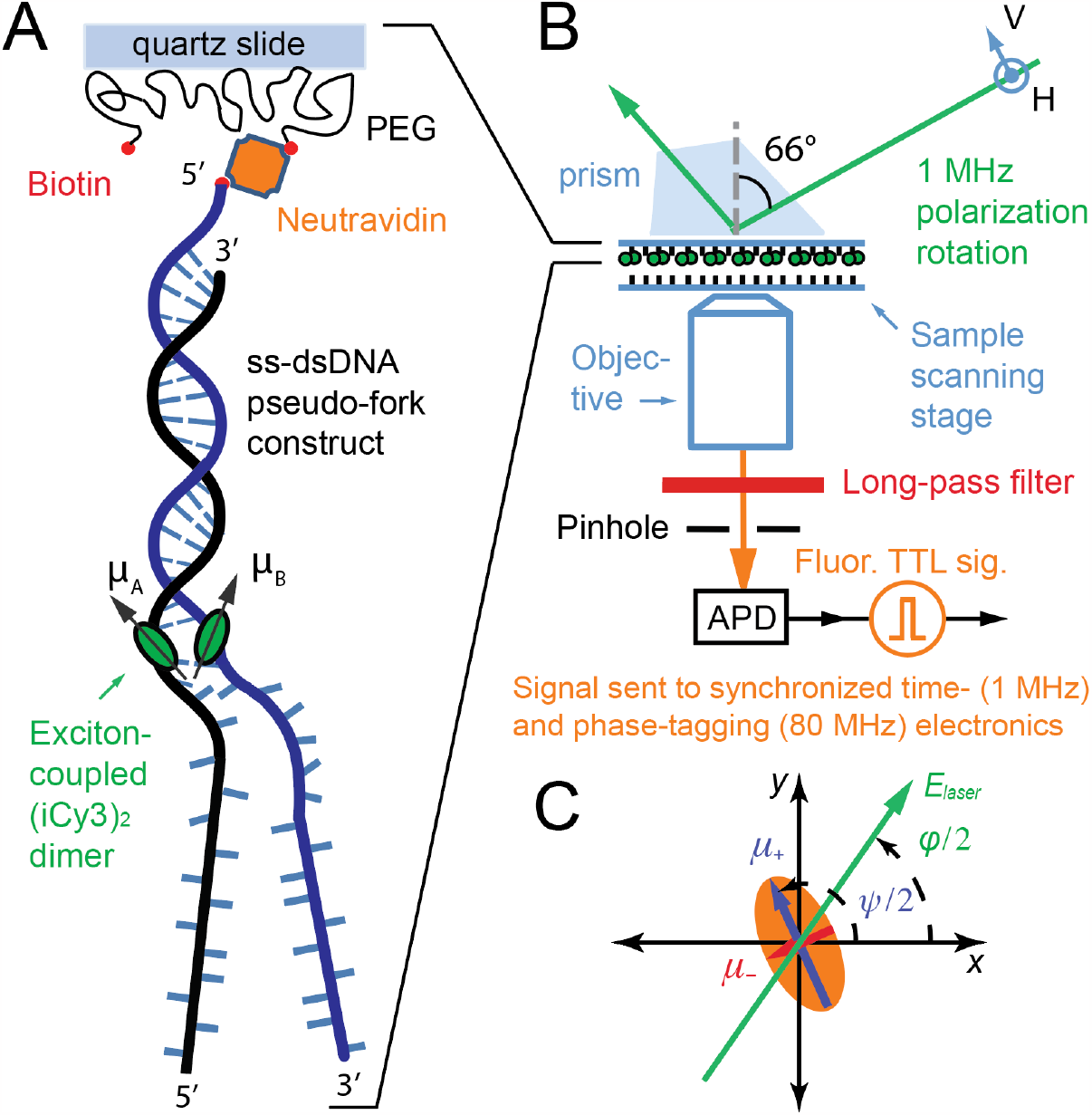
The PS-SMF microscopy experimental layout. (***A***) An (iCy3)_2_ dimer-labeled ss-dsDNA fork construct, which can form a protein-DNA complex, is attached to the microscope slide surface using biotin / neutravidin linkages. (***B***) A total internal reflection fluorescence (TIRF) microscope is used to illuminate the sample with a continuous-wave (cw) 532 nm laser. The plane-polarized electric field vector of the laser is continuously rotated at the frequency Ω/2π = 1 MHz by sweeping the phase of an interferometer (see Fig. A1 of Appendix A). Individual signal photons (fluorescence) from a single molecule are detected using an avalanche photodiode (APD). Each detection event is registered within a time window Δ*t* = 1 μs, and its phase is assigned to one of 80 incremented values (with bin width Δφ = 360°/80 = 4.5°) using the method of phase-tagged photon counting (PTPC) [8]. (***C***) A single (iCy3)_2_ dimer-labeled ss-dsDNA construct has symmetric and anti-symmetric EDTMs (labeled μ_±_, shown as blue and red vectors, respectively), which define the major and minor axes of a ‘polarization ellipse’ in the transverse cross-sectional area of the incident laser beam. The orthogonally polarized symmetric and anti-symmetric excitons are alternately excited as the laser electric field vector (shown in green) rotates in the clockwise direction (with phase φ/2 = Ω*t*/2).

PS-SMF experiments provide both structural and kinetic information about the microscopic configurations of the sugar-phosphate backbones and bases immediately surrounding the (iCy3)_2_ dimer probe. PS-SMF trajectories contain information about the equilibrium distribution of conformational macrostates and their associated rates of interconversion, which are directly related to parameters that characterize the site-specific local free energy landscape (FEL) [9]. From the photon count data stream, we determine the following three ensemble-average functions of the PS-SMF signal: (*i*) the two-point time-correlation function (TCF), 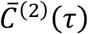 that describes the average correlation between two successive measurements separated by the interval τ, which is sampled on tens-of-microsecond time scales; (*ii*) the three-point TCF,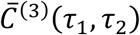that describes the average correlation between three successive measurements separated by the intervals τ_1_ and τ_2_ and is sampled on millisecond time scales; and (*iii*) the probability distribution function (PDF) of the PS-SMF signal visibility *P*(*v*), which is averaged over the 10 ms time scale required to achieve a signal-to-noise (S/N) ratio of ∼10.

Although beyond the scope of the present paper, the above functions can be analyzed using a kinetic network model to characterize the equilibrium and kinetic properties of multistep conformational transition pathways involved in such systems [10]. While the two-point TCF, 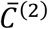, contains information about the characteristic relaxation times of the system, the three-point TCF,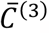, contains additional information about the exchange times between pathway intermediates [11-16].

Recently, Heussman *et al*. used ensemble absorbance, circular dichroism (CD) and two-dimensional fluorescence spectroscopy (2DFS) to study the *ensemble average* local conformations and conformational disorder of iCy3 monomer- and (iCy3)_2_ dimer-probes at various positions within ss-dsDNA fork constructs [5-7]. From these studies it was concluded that the average conformation of the (iCy3)_2_ dimer-probe changes systematically as the labeling probe site is varied across the ss-dsDNA junction from the +2 to the -2 positions (see Fig. 1*D* for site-labeling nomenclature). For positive integer positions the mean local conformation was shown to be right-handed (defined as exhibiting a mean twist angle, 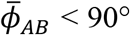), and for negative integer positions the local conformation was left-handed 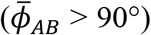. Moreover, 2DFS experiments indicated that the distributions of conformational states at these key positions are relatively narrow [6], suggesting that regions of the junction extending towards and into the single-stranded segments exhibit moderate levels of structural order.

A possible explanation for the results of Heussman *et al*. [6, 7] is that the (iCy3)_2_ dimer probe, which is covalently linked to the sugar-phosphate backbones and bases immediately surrounding the probe, can adopt only a small number of local conformations whose relative stabilities depend on position relative to the ss-dsDNA fork junction. In Fig. 3, we illustrate this concept using a hypothetical free energy landscape, which depicts four possible local conformations of the (iCy3)_2_ dimer probe at a positive integer position, with the right-handed Watson-Crick (WC) B-form conformation being most favored. In this picture the free energy differences between the B-form ground state and the other (non-canonical) local conformations are sufficiently high to ensure that the Boltzmann-weighted distribution of available macrostates is dominated by the B-form structure. At the same time, the moderate free energies of activation allow for infrequent transitions between non-canonical local structures, which are present in trace amounts. As the probe-labeling position is varied across the ss-dsDNA junction towards the single-stranded region, one might expect the free energy surface to exhibit local minima with approximately the same coordinate values, but with relative stabilities and barrier heights shifted to reflect the presence of non-canonical structures observed in ensemble measurements [5-7].

**Figure 3.**
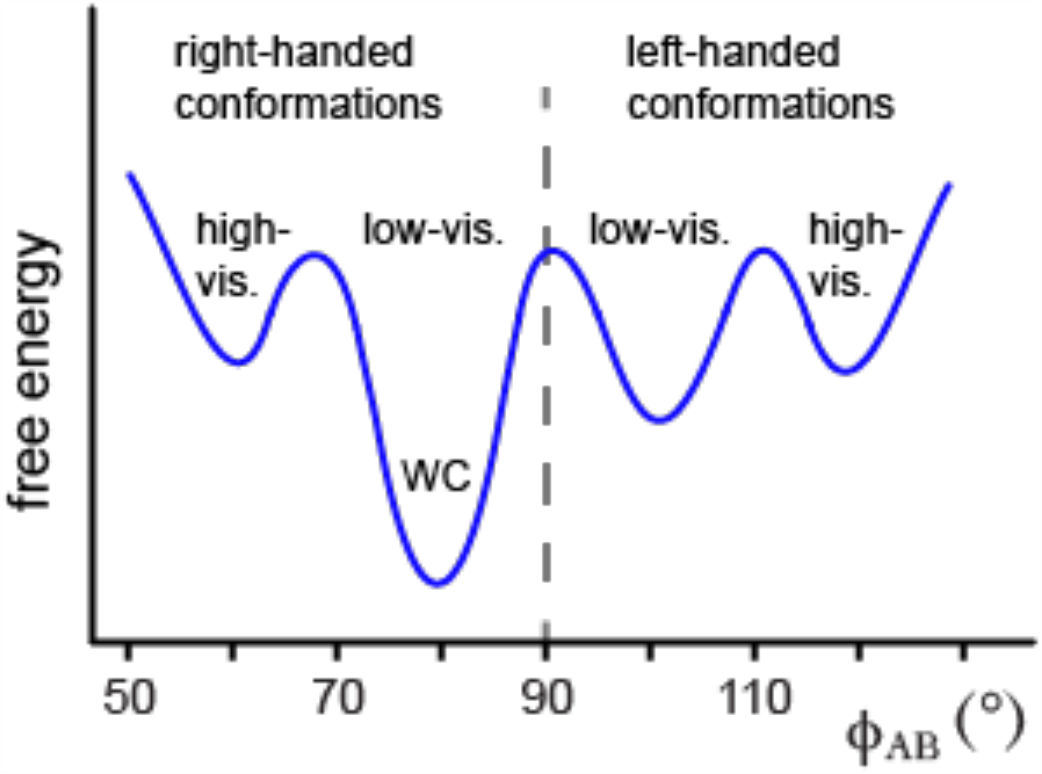
A hypothetical free energy surface (FES) describing local fluctuations of the (iCy3)_2_ dimer-labeled sugar-phosphate backbones near the ss-dsDNA fork junction as a function of the twist angle parameter, *ϕ*_*AB*_ [6].

The PS-SMF method presented here is an extension of one previously developed by Phelps *et al*. [17], which simultaneously monitors the linear dichroism (LD) and Förster resonance energy transfer (FRET) signals from an internally labeled iCy3 / iCy5 donor-acceptor pair. The primary advantage of the PS-SMF method is that it can sensitively isolate signals due to the relative internal motions of an exciton-coupled (iCy3)_2_ dimer probe from those arising from collective rotation and/or excited state population fluctuations (e.g., ‘blinking’) [18]. As such, PS-SMF experiments provide a direct means to observe the dynamic interconversions between local conformational macrostates of (iCy3)_2_ dimer-labeled ss-dsDNA constructs. The PS-SMF method is thus well suited for studies of the kinetic pathways and equilibrium distributions of the conformational fluctuations of the (iCy3)_2_ dimer-labeled sugar-phosphate backbones and bases that are central to understanding — at the molecular level — the different forms of local DNA ‘breathing’ at specified positions relative to ss-dsDNA junctions.

In the sections that follow we show that PS-SMF experiments can be used to study position- and salt-dependent ‘breathing’ fluctuations of the (iCy3)_2_ dimer-labeled ss-dsDNA fork constructs that we use as ‘test substrates’ for the PS-SMF methodology we describe in this paper. Among the significant findings of this work is that the local conformations of these (iCy3)_2_ dimer-labeled ss-dsDNA constructs undergo transient fluctuations between four conformational macrostates, as suggested by the free energy landscape depicted schematically in Fig. 3. Furthermore, at probe labeling positions at and near the ss-dsDNA junction that extend towards the single-stranded region, the local B-form conformation becomes unstable relative to conformations that exist in the duplex at only trace occupancy levels. Altering the salt concentration from approximately ‘physiological’ values [19] (6 mM MgCl_2_, 100 mM NaCl) [20] towards either higher or lower values also destabilizes the B-form conformation, a finding reminiscent of the concept of cold denaturation [21]. While certain aspects of salt concentration-dependent DNA stability and protein-DNA interactions are well understood [22-28], little information is currently available about the microscopic details of how different salt concentrations (here of NaCl and MgCl_2_) perturb conformational fluctuations near ss-dsDNA fork junctions. The results of our experiments suggest a possible structural framework for understanding the roles of transient DNA conformational fluctuations at and near ss-dsDNA fork junctions in driving the processes of protein-DNA complex recognition, assembly, and function.

## 2. Materials and Methods

### 2.1. (iCy3)_2_ dimer-labeled single-stranded (ss) – double-stranded (ds) DNA fork constructs

The model ss-dsDNA fork constructs that we used in this work have either an iCy3 monomer or an (iCy3)_2_ dimer incorporated into the DNA framework, as shown in Figs. 1*A* and 1*B*. The specific nucleic acid base sequences are listed in Table I. We purchased nanomolar quantities of single-stranded oligonucleotides from Integrated DNA Technologies (IDT, Coralville, IA). The initially dehydrated samples were rehydrated at 100 nM concentrations in aqueous buffer (100 mM NaCl, 6 mM MgCl_2_ and 10 mM Tris, pH 8.0). Complementary oligonucleotides were combined in a 1:1.5 molar concentration ratio between the biotinylated and non-biotinylated strands, respectively, before they were heated to 94°C for 4 min and allowed to cool slowly to room temperature (∼23°C). The annealed iCy3 monomer and (iCy3)_2_ dimer-labeled ss-dsDNA constructs contained both single-stranded and double-stranded regions, with the probe labeling positions indicated using the nomenclature described in Fig. 1*D*. The iCy3 monomer-labeled ss-dsDNA constructs contained a thymine base (dT) in the complementary strand at the position directly opposite to the probe chromophore. Solution samples were stored at 4°C between experiments conducted over consecutive days, or frozen at –4°C between experiments conducted over more extended periods.

### 2.2. Sample preparation

Solutions of annealed ss-dsDNA samples were diluted 1000-fold (∼100 pM concentration) before they were introduced into a custom-built microscope sample chamber, as described in prior work [11]. Sample chambers were constructed from microscope slides, which were chemically modified using the procedure of Chandradoss *et al*. [29]. The inner surfaces of the sample chambers were coated with a layer of polyethylene glycol (PEG), which was sparsely labeled with biotin. Neutravidin was used as a linker to bind the biotin-labeled PEG to the biotin molecules covalently attached to the 5’ ends of the dsDNA regions of the ss-dsDNA constructs, as shown in Table 1. To reduce the rate of photobleaching in our measurements, an oxygen scavenging solution consisting of glucose oxidase, catalase and glucose was introduced into the sample cell before the PS-SMF data were recorded. To suppress the ‘photo-blinking’ effects of excited-state intersystem crossing, the triplet state quencher Trolox was used in the solution buffer [18].

**Table 1.**
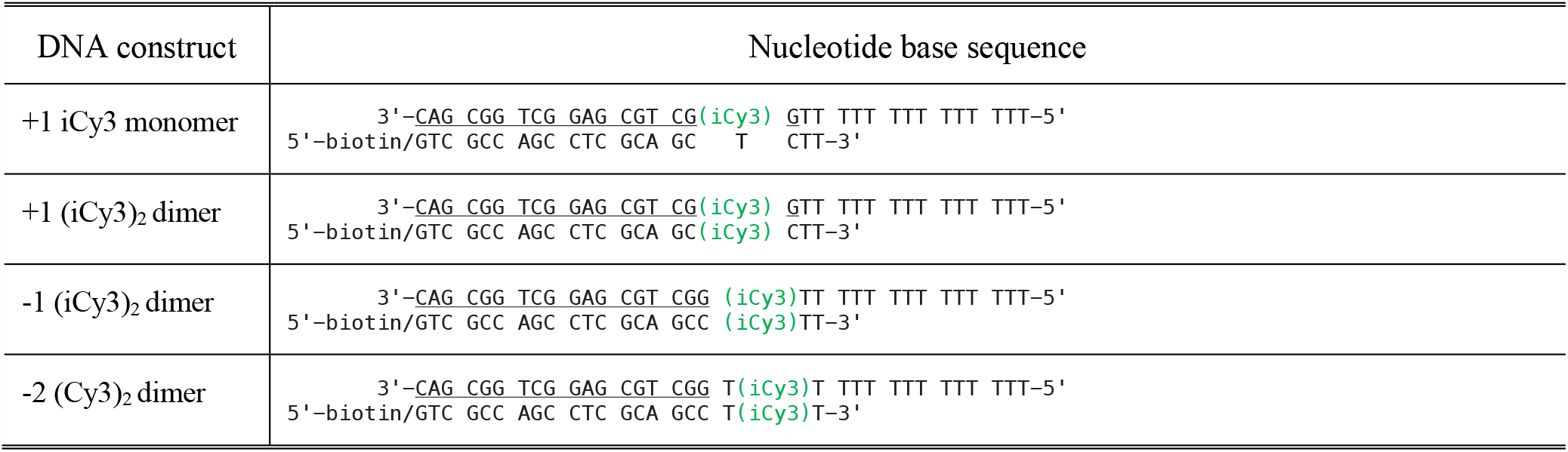
Base sequences and nomenclature for the iCy3 monomer and (iCy3)_2_ dimer-labeled ss-dsDNA fork constructs used in these studies. The horizontal lines indicate regions of complementary base pairing.

### 2.3. Polarization-sweep single-molecule fluorescence (PS-SMF) microscopy

In Fig. 2, we show a schematic of the instrumental setup for PS-SMF experiments on model (iCy3)_2_ dimer-labeled ss-dsDNA fork constructs. The setup is an extension of an approach previously developed to simultaneously monitor single-molecule Förster resonance energy transfer (FRET) and fluorescence-detected linear dichroism (FLD) signals from iCy3/iCy5 (hetero-) dimer-labeled DNA constructs [17]. As we discuss in further detail below, the signals from PS-SMF measurements provide direct information about the distributions of conformational macrostates and the conformational dynamics of the (iCy3)_2_ dimer probes, which are site-specifically positioned within the ss-dsDNA fork junctions.

The (iCy3)_2_ dimer-labeled ss-dsDNA constructs were chemically attached to the surface of a glass microscope slide using biotin/neutravidin linkages, as described in Sect. 2.2 (see Fig. 2*A*). The sample chamber was placed on a computer-controlled translation stage, and the probe-labeled DNA constructs were illuminated, using total internal reflection fluorescence (TIRF) geometry, by a plane-polarized continuous-wave laser (with center wavelength λ_*L*_ = 532 nm, see Fig. 2*B*). The laser beam was focused to a 50 μm diameter spot at the sample, and the incident power was adjusted to ∼15 mW. Fluorescence from the sample was collected using a high numerical aperture objective lens (N.A. = 1.4), and images from single molecules were isolated using a pinhole and a spectral filter (532 nm long-pass), and detected using an avalanche photodiode (APD). The mean total fluorescence photon detection rate (or mean signal flux, 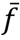) was typically ∼8,000 – 10,000 counts per second (cps). The signal ‘background rate’ was determined to be ∼500 cps by scanning stage positions away from the center of a molecule. Thus, the signal-to-background ratio of our experiments was ∼20. The polarization-dependent signal from each single molecule sample was recorded for a total duration of 30 – 60 s, as described further below.

A Mach-Zehnder interferometer (MZI), equipped with acousto-optic phase modulators in each arm, was used to establish the plane-polarization state of the laser (see Fig. A1 of Appendix A). The relative optical phase of the MZI was swept continuously according to φ = Ω*t*, so that the plane polarization (Jones) vector of the electric field was rotated at the constant frequency Ω/2π = 1 MHz. The electric field of the laser can thus be written (see Appendix B for derivation):

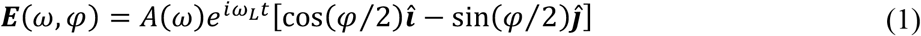

In Eq. (1), *A*(ω) is the (narrow) spectral envelope, ω_*L*_ (=2π*c*/λ_*L*_) is the laser center frequency and 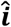 and 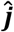 are unit vectors that point along the *x* and *y* axes of the molecular frame, respectively, as shown in Fig. 2*C*. The total electric dipole transition moment (EDTM) of the coupled *AB* dimer is μ_*tot*_ = μ_+_+ μ___, where 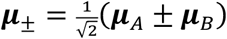 are the EDTMs of the symmetric (+) and anti-symmetric (−) excitons. We note that the magnitudes of the μ_±_ EDTMs depend sensitively on the local conformation of the *AB* dimer (see Fig. 1*C* and discussions in [4-6]). Furthermore, the directions of μ_±_ have orthogonal relative orientation, and thus define a ‘polarization ellipse’ in the cross-sectional area in which the laser beam projects onto the molecular frame:

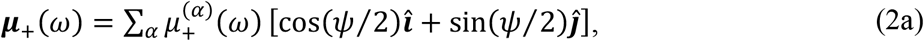

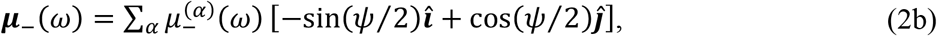

In Eq. (2), the index α enumerates the EDTMs between electronic ground and excited states in order of increasing energy and the angle ψ/2 specifies the orientation of the dimer in the *x*-*y* plane, which we assume to be constant or slowly varying on the time scales of the dimer conformational fluctuations of interest.

At any instant, the ‘polarized’ single-molecule fluorescence intensity is given by the spectral overlap and square modulus of the summed projections between the laser electric field and the total EDTM:

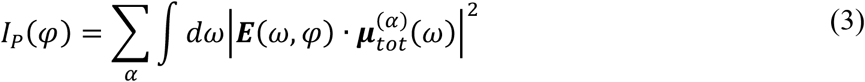

Substitution of Eq. (1) and Eq. (2) into Eq. (3) leads to (see Appendix C for derivation)

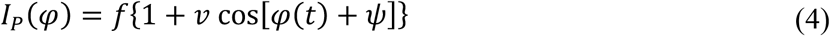

where 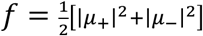 is the signal flux and *v* = [|μ_+_|^2^ − |μ___|^2^]/[|μ_+_|^2^+ |μ___|^2^] is the ‘signal visibility’ with Wμ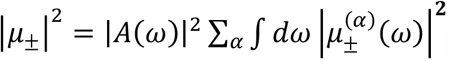

From Eq. (4), we see that the signal is a sum of two terms. The first term is the ‘instantaneous’ signal flux *f*, which is assumed to be constant during a particular measurement period, and the second (φ-dependent) term varies rapidly at the phase modulation frequency (φ = Ω*t*, Ω = 1 MHz). The modulated signal component is proportional to the visibility, *v*, which is directly related to the local conformation of the exciton-coupled (iCy3)_2_ dimer probe. The signal phase ψ represents the orientation of the dimer in the lab frame. Thus, to monitor the internal conformational fluctuations of the (iCy3)_2_ dimer probe, it is necessary to isolate the modulated signal component *v* from the signal flux *f* subject to a well-defined phase ψ.

Under conditions of high signal intensity, a linear detector, such as a photomultiplier tube (PMT), might be used in place of the APD illustrated in Fig. 2*B*. The continuously varying signal described by Eq. (4) could then be measured using a conventional lock-in amplifier [8]. However, under the low-signal-flux conditions of the current work, Eq. (4) describes the probability that the APD detects a single photon at time *t*. Thus, at any instant, we consider each detection event to be a discrete sample of the signal phase given by the Dirac delta function δ(φ − φ_)_), where the φ_)_ parameters are random variables. In our single-molecule experiments, the mean signal flux (integrated over many seconds) was typically 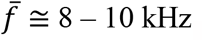, so that on average only one detection event occurred per 100 – 125 modulation cycles. Therefore, to sample the low flux signal as efficiently as possible, we implemented the phase-tagged photon counting (PTPC) method [8]. In this approach, the electronic waveform used to drive the relative phase of the interferometer was synchronized to an 80 MHz digital counter (see Fig. A1 of Appendix A). We thus collected a stream of sparsely sampled single-photon detection events in which we assigned a ‘time stamp,’ *t*_*n*_, with resolution Δ*t* = 1 μs, and a ‘phase stamp,’ φ_*n*_, with resolution Δφ = 360° / 80 = 4.5°, to the *n*th detection event. In post-data-acquisition, we calculated the discrete signal intensity from the set of *N* detected photons that were measured during an (adjustable) sampling window *T*_w_ centered at a particular time *t*.

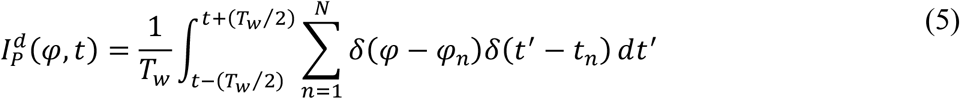

We isolated the modulated component of Eq. (5) by calculating the first term of the complex-valued Fourier series expansion with respect to φ [8].

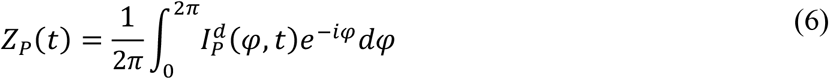

Substitution of Eq. (5) into Eq. (6) shows that the modulated signal component is the sum of *N* single photon phase factors:

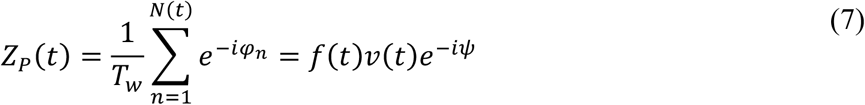

The second equality of Eq. (7) relates the visibility to the modulated signal component according to *v*(*t*) = ⌈*Z*_p_(*t*)⌉/*f*(*t*), where *f*(*t*) = *N*(*t*)/*T*_w_ is the instantaneous signal flux and 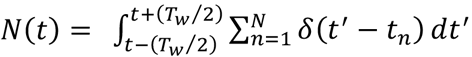 is the number of photons measured at time *t* during the sampling window *T*_w_. We note that the instantaneous flux can be obtained by averaging the signal [Eq. (4)] over the phase variable: 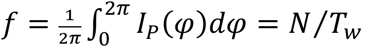.

To demonstrate that the polarization state of the source laser is well described by Eq. (1), and that the phase (φ)-dependent signal behaves according to Eq. (4), we carried out control measurements on: (*i*) an isotropic solution of Rhodamine B in ethanol, and (*ii*) an anisotropically-stretched poly vinyl alcohol (PVA) film containing Cy3 (see Fig. S1 of the SI). As expected, the fluorescence signal of the anisotropic sample exhibited a pronounced φ-dependent modulation, while no significant φ-dependent modulation was observed from the isotropic sample.

### 2.4 Characterization of conformational dynamics using multi-point time-correlation functions (TCFs)

We monitored the local conformational fluctuations of the (iCy3)_2_ dimer-labeled ss-dsDNA constructs as reflected by measurements of the PS-SMF signal visibility, *v*(*t*). We analyzed these fluctuation data by constructing ensemble averaged probability distribution functions (PDFs) and multi-point time-correlation functions (TCFs). In the current work, we provide a qualitative interpretation of our results. However, by applying a kinetic network model analysis like the approach of Israels *et al*. [11], we may obtain from these data equilibrium distributions of macrostates and kinetic rate constants necessary to parameterize the free energy landscapes of these systems. A quantitative analysis of the free energy landscapes is the focus of a forthcoming paper [10].

We characterized the kinetic information contained in our PS-SMF experiments using two-point and three-point TCFs [11-16, 30]. We define the time-dependent fluctuation of the visibility δ*v*(*t*) = *v*(*t*) − ⟨*v*⟩, where ⟨*v*⟩ is the mean value determined from a time series of measurements performed on a single molecule. The two-point TCF is the average product of two successive measurements separated by the interval τ:

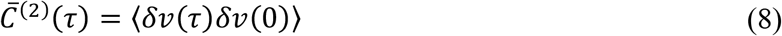

In Eq. (8), the angle brackets indicate a running average over all possible initial measurement times. The function 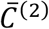 contains information about the number of quasi-stable macrostates of the system, the mean visibility of each macrostate, and the characteristic time scales of interconversion between macrostates [11-13].

The three-point TCF is the time-averaged product of three consecutive measurements separated by the intervals τ_1_ and τ_2_.

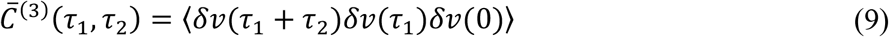

The function 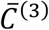 contains detailed kinetic information about the transition pathways that connect the macrostates of the system. This function is sensitive to the roles of intermediates, whose presence can facilitate or hinder successive transitions between macrostates [11]. Unlike even-moment TCFs, the three-point TCF,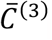, contains no underlying noise-related background term that could potentially obscure the experimentally derived surface [16].

In our analysis, we constructed from our time-dependent measurements of the PS-SMF visibility the PDF, *P*(*v*), using the sampling window *T*_w_ = 10 ms. We chose this value for *T*_w_ because (as shown below) it is roughly equal to the time required for the signal-to-noise ratio (S/N) to acquire a value of ∼10. In addition, we calculated two-point and three-point TCFs, 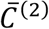 and 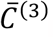, respectively, using the sampling window *T*_w_ = 250 μs. Typically, ∼150 to 250 individual single-molecule data sets were combined to construct each of the experimentally derived statistical functions.

## 3. Results and Discussion

### 3.1 Analysis of PS-SMF spectroscopic signals

For each of the iCy3 monomer and (iCy3)_2_ dimer-labeled ss-dsDNA constructs that we studied (see Table 1) we recorded PS-SMF data streams for a typical duration of ∼30 – 50 s. In Fig. 4, we present examples of PS-SMF signal trajectories for the +1 (iCy3)_2_ dimer-(see Figs. 4*A* – 4*C*) and the +1 iCy3 monomer-labeled ss-dsDNA fork construct (Figs. 4*D* – 4*F*) using the integration window *T*_w_ = 100 ms. In these examples, the mean flux is 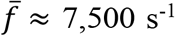 for the (iCy3)_2_ dimer and 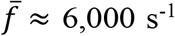 for the iCy3 monomer. We note that the apparent ‘noise’ in the signal trajectories is due to the measurement uncertainty associated with the finite number of photons detected within the integration window at these low flux levels [8]. Thus, the mean signal-to-noise ratio is 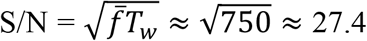 for the trajectory of the (iCy3)_2_ dimer-labeled construct, and S/N ≈ 24.5 for the trajectory of the iCy3 monomer-labeled construct.

**Figure 4.**
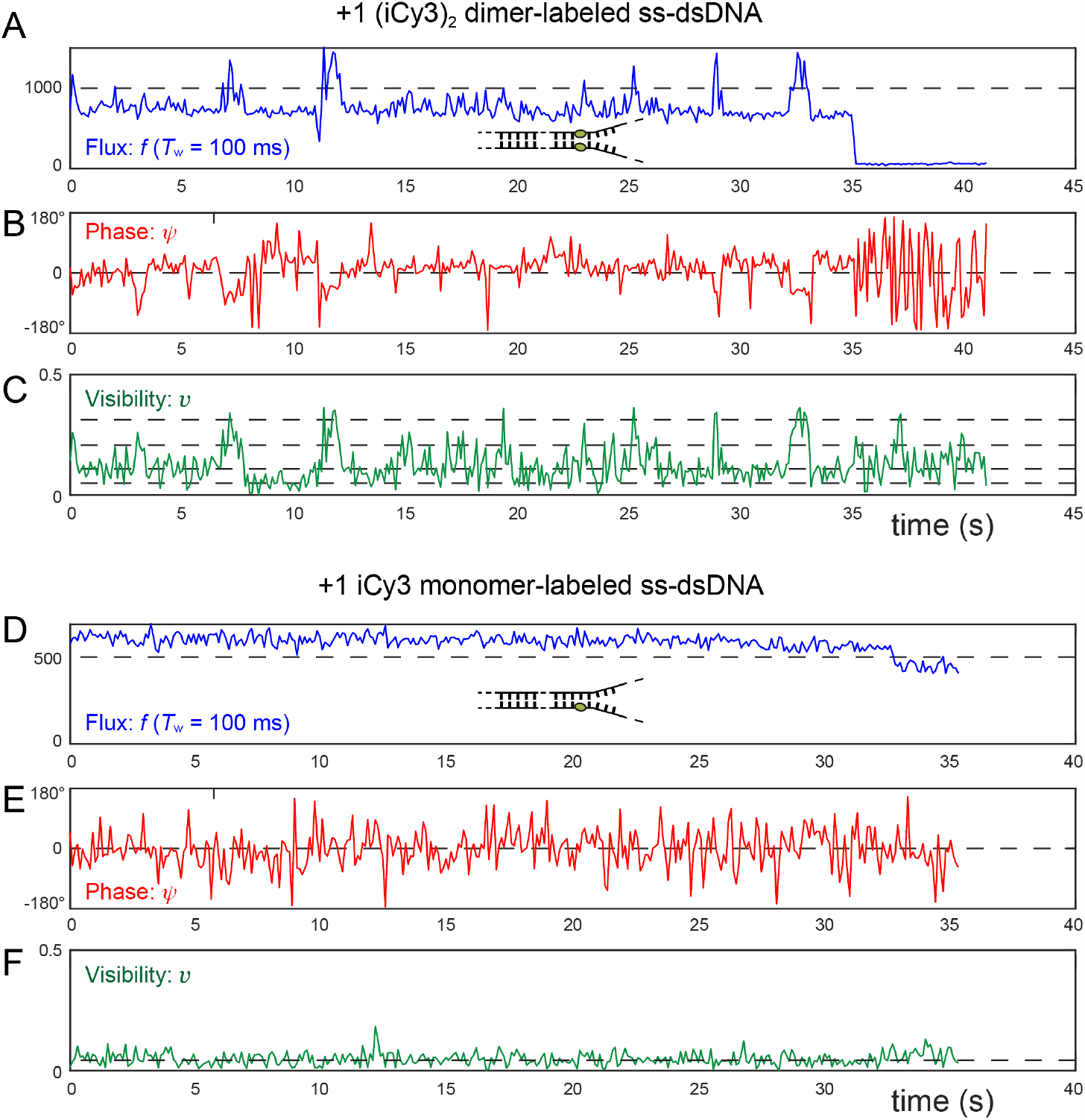
Example PS-SMF signal trajectories using the integration window *T*_*w*_ = 100 ms for the (panels ***A*** – ***C***) +1 (iCy3)_2_ dimer- and (panels ***D*** – ***F***) +1 iCy3 monomer-labeled ss-dsDNA fork constructs. Experiments were performed using 100 mM NaCl and 6 mM MgCl salt concentrations. For the +1 (iCy3)_2_ dimer-labeled construct the photon data stream was recorded with a mean signal flux 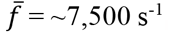, corresponding to S/N ≈ 27.4. For the +1 iCy3 monomer-labeled construct the mean signal flux was 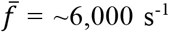, corresponding to S/N ≈ 24.5. In panels ***A*** and ***D***, the instantaneous signal flux, *f*(*t*) = *N*(*t*)/*T*_*w*_, is shown in blue. In panels ***B*** and ***E***, the instantaneous signal phase, ψ, is shown in red, ‘unwrapped’ with its mean value set equal to zero, 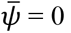. In panels ***C*** and ***F***, the signal visibility, *v*(*t*), is plotted in green. Horizontal dashed lines are shown as a guide to the eye to indicate the presence of multiple discrete conformational states for the (iCy3)_2_ dimer-labeled ss-dsDNA construct (panel ***C***), while only a single discrete state is observed for the iCy3 monomer-labeled ss-dsDNA construct (panel ***F***).

The three signal components are the instantaneous flux, *f*(*t*) = *N*(*t*)/*T*_w_ (Figs. 4*A* and 4*D*), the phase, ψ(*t*) (Figs. 4*B* and 4*E*) and the visibility, *v*(*t*) = ⌈*Z*_P_(*t*)⌉/*f*(*t*) (Figs. 4*C* and 4*F*), which are described by Eqs. (5) – (7), respectively. The flux, *f*(*t*), is equivalent to the fluorescence intensity, a quantity that is independent of the laser polarization since it is averaged over the polarization phase variable. We note that fluctuations of the flux reflect changes in the local environment of the (iCy3)_2_ dimer probe, which influence excited state deactivation pathways. The phase, ψ(*t*), is shown ‘unwrapped’ with its mean value set (arbitrarily) to 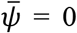. The phase represents the orientation of the major axis of the (iCy3)_2_ dimer ‘polarization ellipse’ in the laboratory frame (see Fig. 2*C*), which undergoes intermittent jumps about its mean value 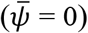 over the duration of the trajectory. For the dimer, a phase jump on the order of 180° is expected to occur when the inter-molecular twist angle *ϕ*_*AB*_ (see Fig. 1*C*) undergoes an abrupt fluctuation traversing the value *ϕ*_*AB*_ = 90°, which leads to a juxtaposition of the symmetric and anti-symmetric excitons. For both the monomer and the dimer, the phase jumps appear to occur randomly over the full -180° to 180° range, which we attribute to uncertainty associated with low signal sampling and other noise sources such as room vibrations.

From Fig. 4*C*, we see that the visibility trajectory for the +1 (iCy3)_2_ dimer-labeled ss-dsDNA construct undergoes discontinuous transitions between a small number of discrete values within the range 0 < *v* < 0.4 (dashed lines are guides to the eye). In this case, the visibility is a direct measure of the internal conformation changes of the (iCy3)_2_ dimer probe, as discussed in Sect. 2.3 [see Eq. (4)]. In contrast, the visibility trajectory for the +1 iCy3 monomer-labeled ss-dsDNA construct shown in Fig. 4*F* exhibits relatively uniform fluctuations about a single discrete value, *v* ∼0.08. Here, the visibility measures only the mean projection of the iCy3 monomer’s ‘polarization ellipse’ within the transverse plane of the polarized laser beam (Fig. 2*C*), which remains relatively constant over the duration of the trajectory. For example, were the EDTM of the iCy3 monomer to be oriented orthogonally to the laser polarization, the visibility would be zero. In principle, the visibility can assume values within the continuous range -1 < *v* < +1, with sign inversion corresponding to a 180° phase shift. Nevertheless, the PS-SMF experiment cannot distinguish between negative and positive values of the visibility, so that in practice we measure the absolute value |*v*|.

From the PS-SMF data streams, we constructed probability distribution functions (PDFs) for different values of the integration time window, *T*_*w*_. In Fig. 5, we compare PDFs of the flux, *f* (left column), visibility, *v* (middle), and phase, ψ (right), for the +1 (iCy3)_2_ dimer-labeled ss-dsDNA construct. Additional PDFs for the +1 (iCy3)_2_ dimer- and the +1 iCy3 monomer-labeled ss-dsDNA constructs using *T*_*w*_ = 500 ms are shown in Figs. S2 and S3 of the SI, respectively. For all the samples that we studied, the visibility PDFs are asymmetrically distributed and bounded within the range 0 < *v* < 0.5, with mean value 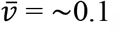. The widths of the visibility PDFs narrow with increasing *T*_*w*_, revealing the presence of distinct features. The effect of increasing *T*_*w*_ is to average over random noise associated with the sparsely detected (photon counting) signal. However, choosing too large an integration period can potentially lead to additional, undesirable narrowing of the PDFs by averaging over the state-to-state interconversion events that we wish to monitor and study [16]. It is therefore important to choose a value of *T*_*w*_ that is large enough to ensure that S/N ≳ 10, but small enough to retain the necessary time resolution to monitor the relevant kinetics. Since the mean flux in our experiments is typically 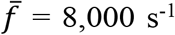, a suitable integration period is *T*_*w*_ = 10 ms, which corresponds to 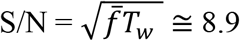. However, we find *T*_*w*_ = 100 ms to be useful for display purposes.

**Figure 5.**
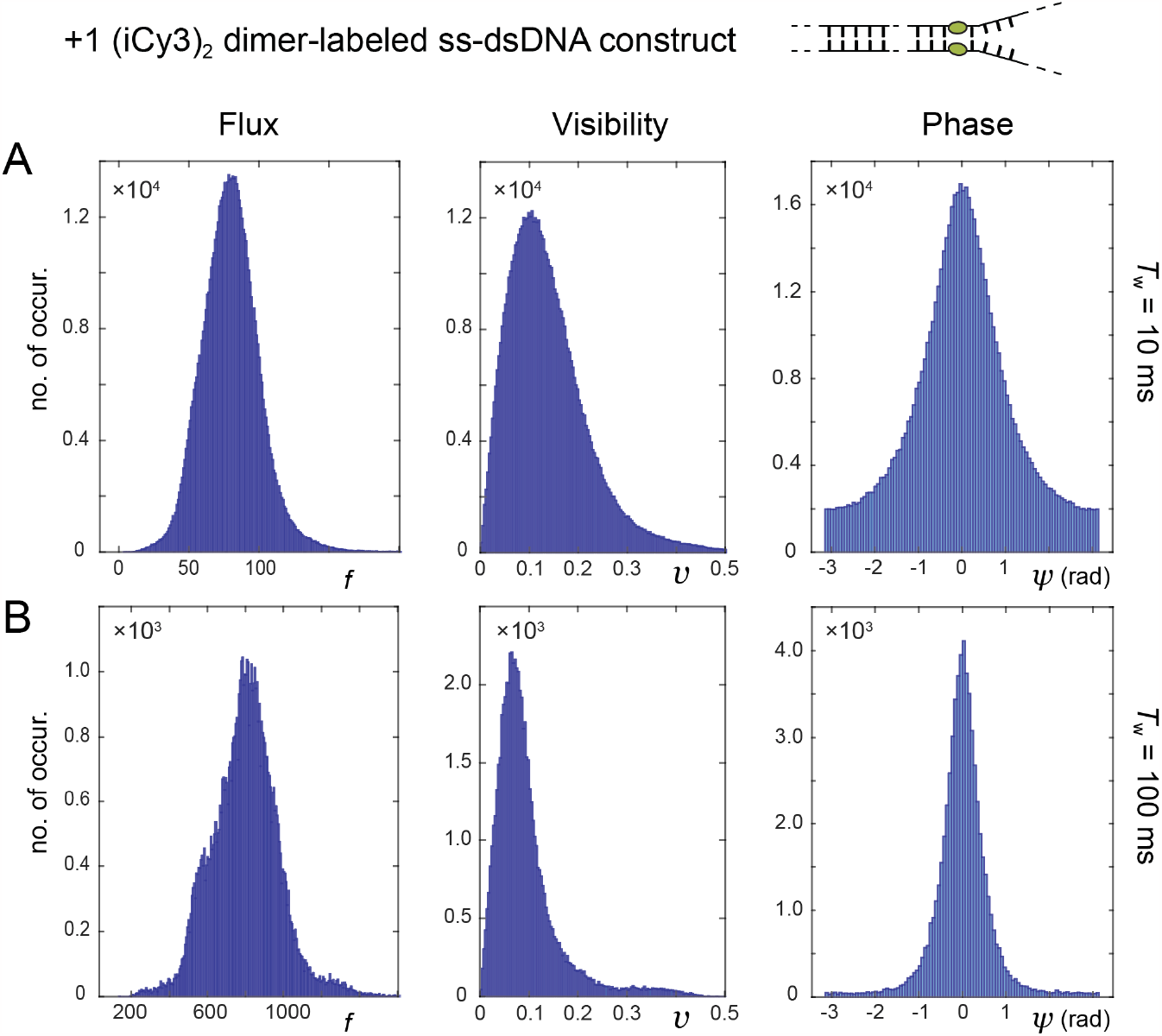
PDFs of the flux (left column), visibility (middle) and phase (right) for the +1 (iCy3)_2_ dimer-labeled ss-dsDNA fork construct. PDFs were constructed from the raw photon data streams (see Fig. 4). The mean signal flux is 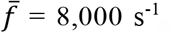. Comparisons are shown varying the integration window (panel ***A***) *T*_*w*_ = 10 ms and (panel ***B***) *T*_*w*_ = 100 ms. The corresponding 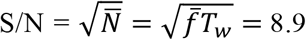 and 28, respectively.

Like the visibility, the phase PDFs narrow with increasing *T*_*w*_, indicating a singular value of the phase that defines the visibility over the integration time window (Fig. 5, right column). However, the flux PDFs do not converge to well-behaved distributions, but rather exhibit increasingly complex structure with increasing *T*_*w*_ (Fig. 5, left column). While the instantaneous flux depends on multiple environmental factors, which influence chromophore absorbance and fluorescence efficiency (e.g., probe orientation and excited electronic state dynamics), the normalized visibility depends primarily on the relative strengths and orientations of the probe EDTMs.

In Fig. 6, we compare the visibility PDF of the +1 (iCy3)_2_ dimer-labeled construct to the +1 iCy3 monomer-labeled construct. For the +1 monomer, the PDF exhibits a single ‘low visibility’ feature in the range 0 ≲ *v* ≲ 0.1 (see Figs. 6*B* and 6*D*), which results from the projection of the monomer EDTM onto the plane of the rotating laser polarization. This contrasts with the +1 dimer-labeled construct, which exhibits – in addition to a dominant low visibility feature – a sparsely populated ‘high visibility’ feature in the range 0.1 ≲ *v* ≲ 0.4 (see Figs. 6*A* and 6*C*). This underlying structure of the visibility PDF for the dimer is reinforced when the integration window is increased to *T*_*w*_ = 500 ms, while that of the monomer exhibits only the single low visibility feature (see Figs. S2 and S3 of the SI). The broadly distributed features that we observe for the (iCy3)_2_ dimer-labeled ss-dsDNA constructs indicate the presence of both stable and thermally activated local conformations of the dimer probes. Transitions between low and high visibility macrostates are evident in the trajectories for the (iCy3)_2_ dimer-labeled ss-dsDNA construct, as shown in Fig. 4*C*, but not for the iCy3 monomer-labeled ss-dsDNA construct (Fig. 4*F*).

**Figure 6.**
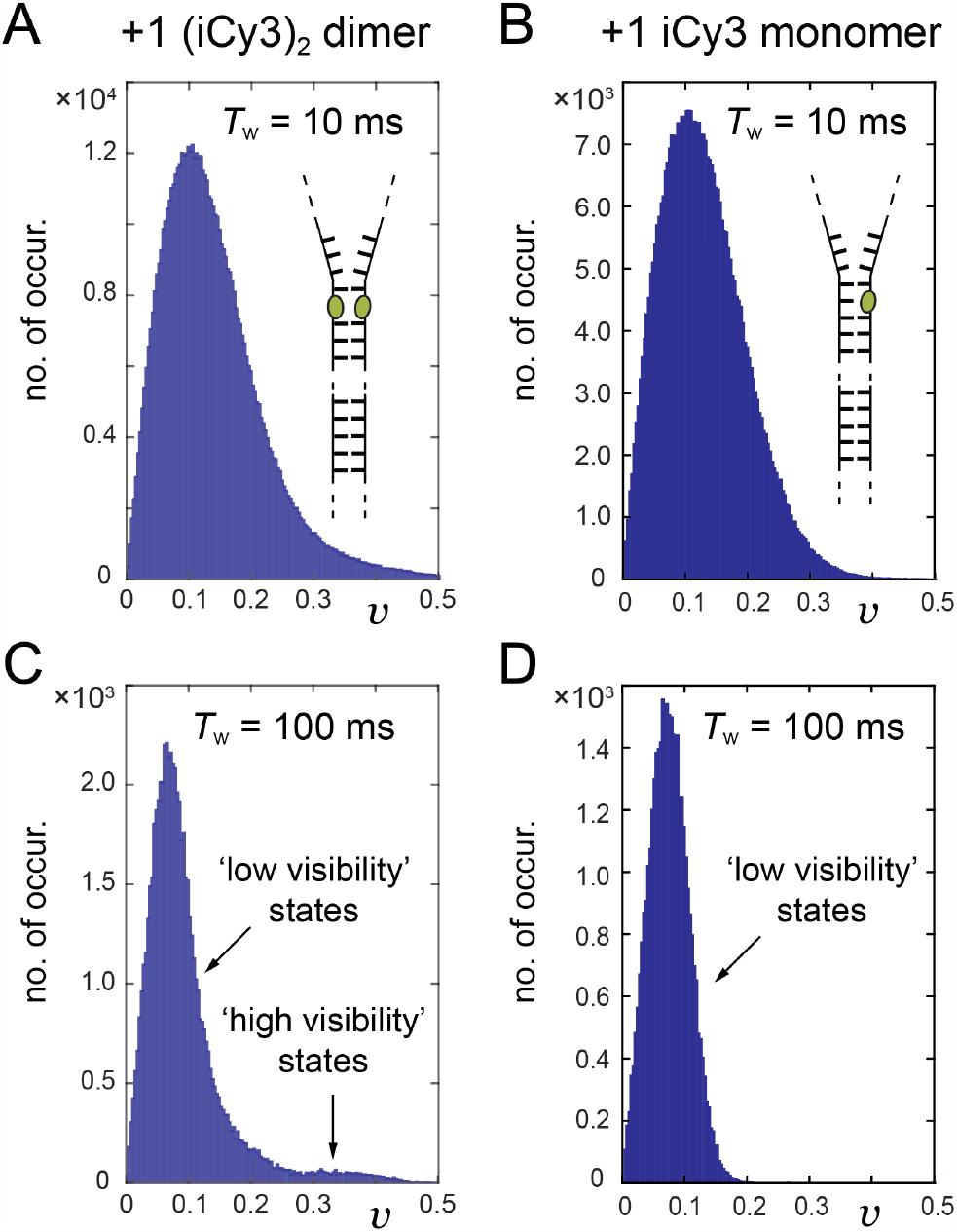
Probability distribution functions (PDFs) of the visibility for (***A, C***) the +1 (iCy3)_2_ dimer-labeled ss-dsDNA construct, and (***B, E***) the +1 iCy3 monomer-labeled ss-dsDNA construct. (***A, B***) *T*_*w*_ = 10 ms. (***C, D***) *T*_*w*_ = 100 ms.

The above observations suggest that the (iCy3)_2_ dimer probes interconvert between a small number of local conformations, which likely depend on the relative stabilities and dynamics of the bases and sugar-phosphate backbones immediately adjacent to the (iCy3)_2_ dimer probes. Furthermore, these results agree with recent ensemble studies of (iCy3)_2_ dimer-labeled ss-dsDNA constructs, which concluded that only a small number of local conformational macrostates of the (iCy3)_2_ dimer probes inserted rigidly into the ss-dsDNA constructs are populated under room temperature and physiological salt conditions due to steric restrictions of the surrounding nucleic acid bases and sugar-phosphate backbones [6].

To determine whether a unique correspondence exists between the instantaneous flux and visibility, we examined the bivariate PDFs for the +1 (iCy3)_2_ dimer- and the +1 iCy3 monomer-labeled ss-dsDNA constructs. In Fig. 7, we plot the bivariate distributions for these constructs as two-dimensional contour diagrams. The bivariate PDF for the +1 (iCy3)_2_ dimer-labeled ss-dsDNA construct (Fig. 7*A*) shows that the low visibility conformations (with 0 ≲ *v* ≲ 0.1) correspond to a broad range of flux values, while the high visibility states (with 0.1 ≲ *v* ≲ 0.4) correspond to relatively low flux values. Similarly, the low visibility states of the iCy3 monomer-labeled ss-dsDNA construct (Fig. 7*B*) correspond to a broad range of flux values. Thus, there does not appear to be a unique (one-to-one) correspondence between flux and visibility values [31], which suggests that only the visibility distribution histograms, but not those for the flux, can be interpreted straightforwardly in terms of the local conformational fluctuations of the iCy3 monomer and (iCy3)_2_ dimer probes.

**Figure 7.**
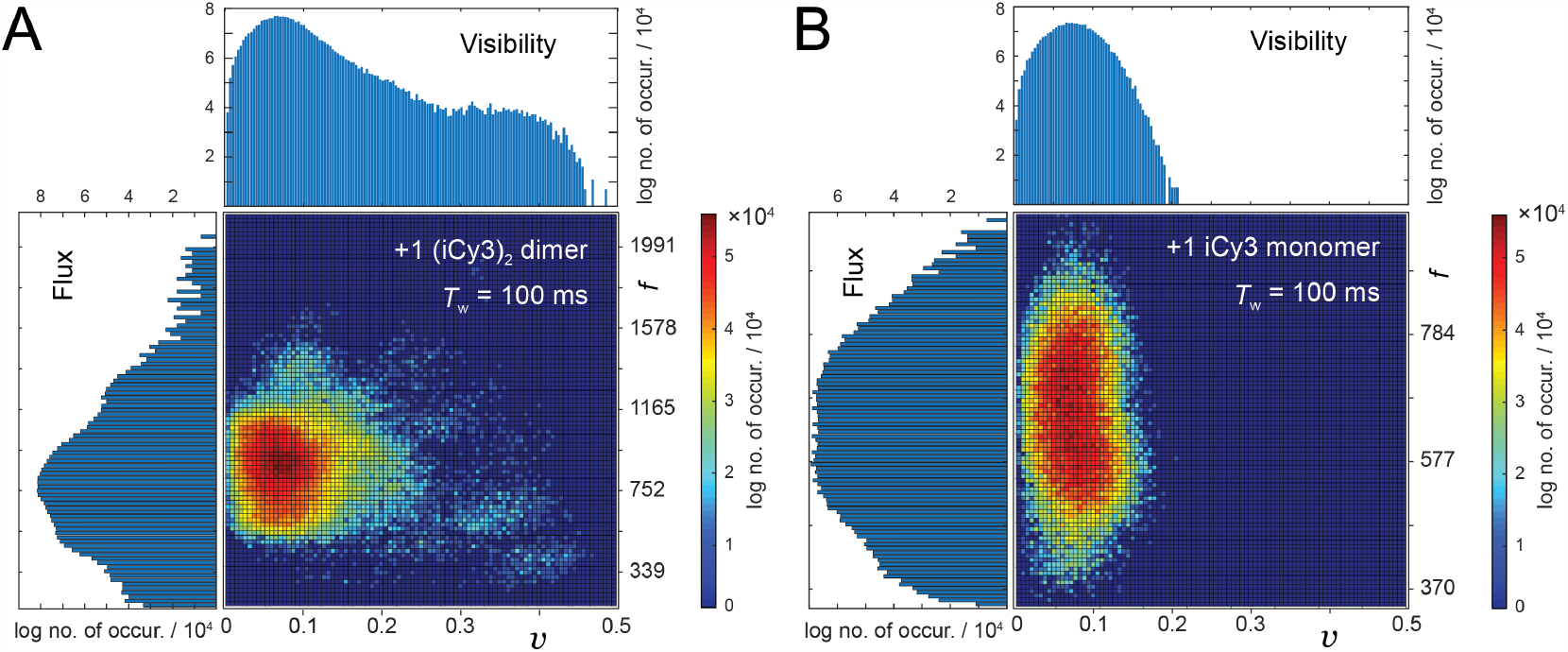
Joint PDFs for the flux and visibility signals (panel ***A***) the +1 (iCy3)_2_ dimer-labeled ss-dsDNA construct and (panel ***B***) the +1 iCy3 monomer-labeled ss-dsDNA construct using *T*_*w*_ = 100 ms.

We next examined how the PS-SMF signals report on the dynamics of the site-specifically-labeled positions within ss-dsDNA fork constructs. In Fig. 8, we plot separately the two-point TCFs for the visibility and the flux distributions for the +1 iCy3 monomer- and the +1 (iCy3)_2_ dimer-labeled ss-dsDNA constructs. For these TCF calculations, we resolved sub-millisecond time scale dynamics by using values for *T*_*w*_ much shorter than those used for the PDF calculations discussed above. The TCFs shown in Fig. 8 use *T*_*w*_ = 250 μs, and these data are plotted alongside model fits to multi-exponential functions, ∑_i_αi exp(− τ/*t*_i_), where the number of decay components for the dimer is 4 and the number of decay components for the monomer is 3. The values that we obtained for the fitting parameters are given in Table 2. Although the time dependencies of the visibility and flux TCFs are qualitatively similar, we found that the decay components of the flux TCFs are generally faster than those of the visibility for all the ss-dsDNA constructs that we studied. Thus, although the dynamic processes that give rise to fluctuations of the flux and visibility signals are related to one another, they are clearly not identical.

**Table 2.** Multi-exponential fit parameters for the flux and visibility 2-point TCFs of the iCy3 monomer-and (iCy3)_2_ dimer-labeled ss-dsDNA constructs. Note that the TCFs and fits are normalized to the point τ = 250 μs. Error bars are +/− one significant figure of the values reported.

| DNA construct | $A_1$ | $t_1 (\mu\text{s})$ | $A_2$ | $t_2 (\text{ms})$ | $A_3$ | $t_3 (\text{ms})$ | $A_4$ | $t_4 (\text{s})$ |
| --- | --- | --- | --- | --- | --- | --- | --- | --- |
| visibility two-point TCFs |  |  |  |  |  |  |  |  |
| +1 (iCy3)<sub>2</sub> dimer | 0.8 | 275 | 0.18 | 2.9 | 0.09 | 43.8 | 0.31 | 2.5 |
| +1 iCy3 monomer | 0.98 | 493 | 0.32 | 4.2 | 0.03 | 1,200 | — | — |
| flux two-point TCFs |  |  |  |  |  |  |  |  |
| +1 (iCy3)<sub>2</sub> dimer | 0.92 | 265 | 0.17 | 3.9 | 0.09 | 56.7 | 0.32 | 2.7 |
| +1 iCy3 monomer | 0.99 | 373 | 0.45 | 3.0 | 0.03 | 1,300 | — | — |

**Figure 8.**
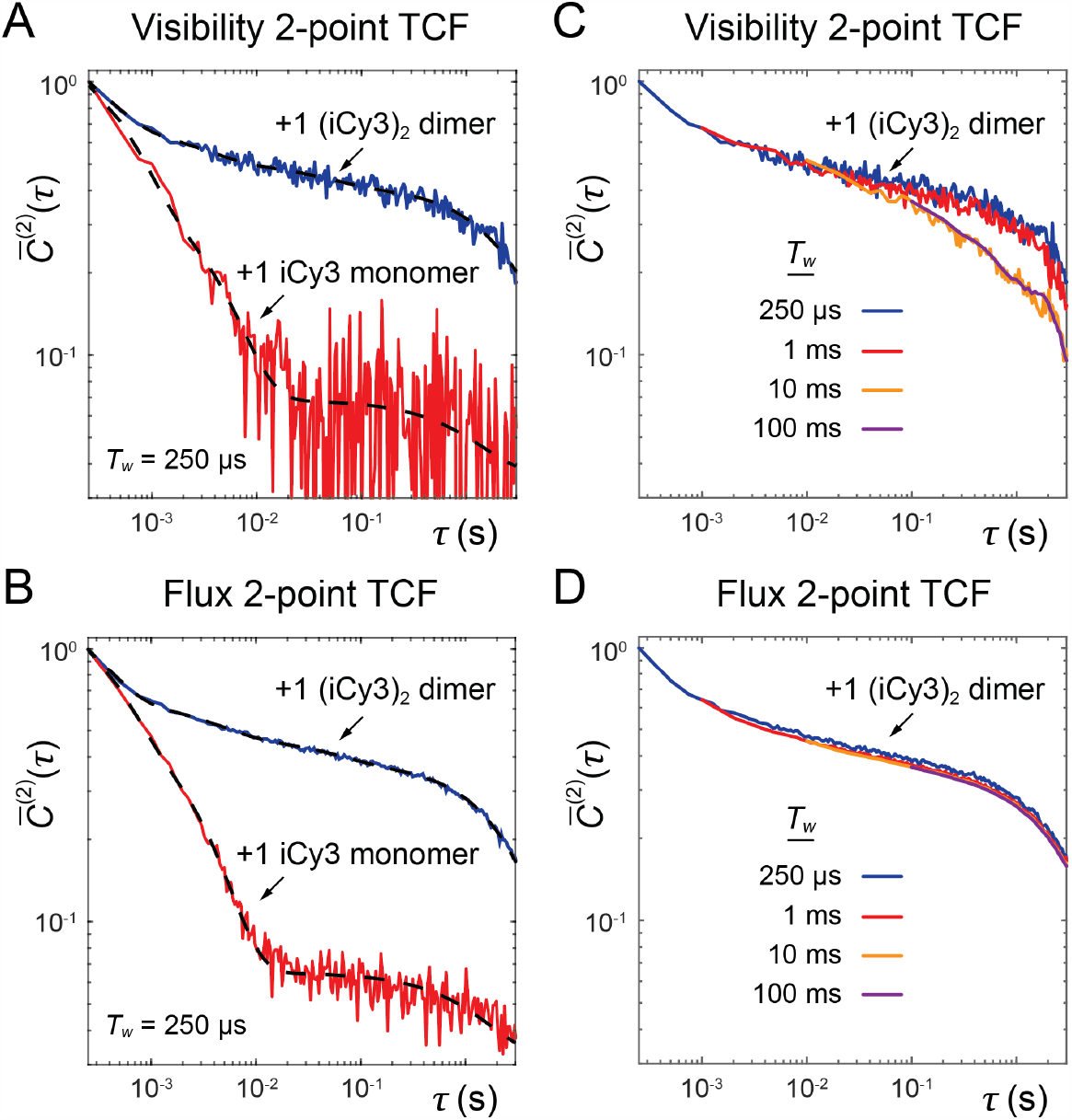
Two-point time correlation functions (TCFs) of (panels ***A*** and ***C***) the visibility, ⟨δ*v*(τ)δ*v*(0)⟩, and (panels ***B*** and ***D***) the flux, ⟨δ*f*(τ)δ*f*(0)⟩, for the +1 iCy3 monomer-labeled ss-dsDNA construct and the +1 (iCy3)_2_ dimer-labeled ss-dsDNA construct. The visibility (panel ***A***) and the flux (panel ***B***) TCFs, respectively, are shown using the integration time window *T*_*w*_ = 250 μs. Solid dashed curves are multi-exponential fits to the data with fitting parameters given in Table 2. The visibility (panel ***C***) and the flux (panel ***D***) TCFs, respectively, of the (iCy3)_2_ dimer-labeled ss-dsDNA construct are shown as a function of *T*_*w*_. Note that all TCFs and fits are shown normalized to the point τ = 250 μs.

From Figs. 8*A* and 8*B*, we see that the decay of the +1 iCy3 monomer-labeled ss-dsDNA construct is significantly faster than the +1 (iCy3)_2_ dimer-labeled construct. The decay of the monomer-labeled construct is dominated by two relatively fast components (*t*_1_ ≲ 0.5 ms, *t*_*2*_ ≲ 4 ms), and exhibits a slowly-decaying, low amplitude, ‘baseline’ with *t*_3_≲1.2 s and amplitude *A*_3_ ∼0.03. In comparison, the (iCy3)_2_ dimer-labeled construct exhibits much slower relaxation behavior with four well-separated decay components: *t*_1_ ∼0.28 ms, *t*_*2*_ ∼2.9 ms, *t*_3_ ∼44 ms and *t*_4_ ∼2.5 s. We note that the slowest time scale component (on the order of seconds) is present in all our data sets. This slow component is also present in our control measurements, which were performed on Cy3 monomer in stretched PVA films (see Fig. S4 of the SI). We thus attribute this slow, low-amplitude process to room vibrations.

Our finding that the two-point TCF for the +1 iCy3 monomer-labeled ss-dsDNA construct exhibits a relatively fast decay is consistent with ensemble spectroscopic measurements [6, 9], from which we concluded that the local environment immediately surrounding the iCy3 monomer probe is structurally disordered. We speculated that the disorder is due to misalignment of complementary Watson-Crick base pairing, which is induced by the presence of the iCy3 monomer in one strand and a thymidine spacer at the opposite position in the complementary strand. In contrast, the relatively slow decay of the +1 (iCy3)_2_ dimer-labeled ss-dsDNA construct suggests that the local environment immediately surrounding the (iCy3)_2_ dimer probes is relatively well ordered at room temperature and under physiological buffer salt conditions, which is also in agreement with results of our prior ensemble studies [6].

We next considered the effects of varying the integration window, *T*_*w*_, on the TCFs of the +1 (iCy3)_2_ dimer-labeled ss-dsDNA construct (see Figs. 8*C* and 8*D*). On the shortest time scale (0.25 ms < τ < 10 ms), the flux and visibility TCFs exhibit similar decays for all values of *T*_*w*_. However, for larger values of *T*_*w*_ (= 10 ms, 100 ms), the decay of the visibility TCFs at longer times (10 ms < τ < 100 ms) is faster than those using small values of *T*_*w*_ (= 250 μs, 1 ms, see Fig. 8*C*). In contrast, the long-time decay behavior of the flux TCFs do not vary with *T*_*w*_ (Fig. 8*D*). Evidently, the relatively slow conformational motions of the (iCy3)_2_ dimer-labeled ss-dsDNA constructs that occur on tens-of-milliseconds and longer time scales, and which are reflected by fluctuations of the visibility, require larger values of *T*_*w*_ to be represented accurately by the TCFs. Thus, the conformational dynamics of the (iCy3)_2_ dimer on time scales less than 10 ms appear to be reflected by fluctuations of both the flux and visibility signals, while the dynamics on time scales greater than 10 ms are most accurately reflected by fluctuations of the visibility alone. In the calculations that follow, we constructed two-point TCFs of the visibility by stitching together the decays using *T*_*w*_ = 250 μs over the range 250 μs – 25 ms, and using *T*_*w*_ = 10 ms over the range 25 ms – 2.5 s.

### 3.2 Studies of local DNA breathing of +1, -1 and -2 (iCy3)_2_ dimer-labeled ss-dsDNA fork constructs

Having established protocols for constructing experimentally derived functions of the visibility (i.e., the PDFs, and the two-point and three-point TCFs), we next used our series of replication fork constructs to examine the sensitivity of these functions to varying (iCy3)_2_ dimer probe positions relative to the ss-dsDNA fork junction. In Figs. 9*A* – 9*C*, we present ensemble results for the +1, -1 and -2 (iCy3)_2_ dimer-labeled ss-dsDNA constructs, which were obtained at room temperature (23°C) and physiological buffer salt conditions ([NaCl] = 100 mM, [MgCl_2_] = 6 mM). The left column shows the visibility PDFs, *P*(*v*), the middle column shows the two-point TCFs, 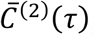, which are overlaid with multi-exponential fits (parameters listed in Table 3), and the right column shows contour plots of the three-point TCFs,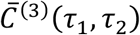. Like the PDF of the +1 (iCy3)_2_ dimer-labeled ss-dsDNA construct (discussed above and shown in Fig. 8*A*), the PDFs of the -1 and -2 dimer-labeled constructs exhibit a major ‘low visibility’ feature in the region 0 ≲ *v* ≲ 0.1, and minor ‘high visibility’ features in the region 0.1 ≲ *v* ≲ 0.4. We find that the relative weights of the ‘high visibility’ features depend on (iCy3)_2_ dimer probe labeling position relative to the ss-dsDNA fork junction, with ‘high visibility’ features notably less prominent for the -1 construct than for the +1 and -2 constructs. Furthermore, the PDF of the -2 construct is significantly different from that of the +1 construct, as expected given the differential stabilities of stacked bases within dsDNA versus ssDNA regions spanning the ss-dsDNA fork junction.

**Table 3.** Multi-exponential fit parameters of the visibility 2-point TCFs for the (iCy3)_2_ dimer-labeled ss-dsDNA fork constructs studied in this work. Note that the TCFs and fits are normalized to the point τ = 250 μs. Error bars are +/− one significant figure of the values reported.

| Construct and salt conditions | $\alpha_1$ | $t_1$ ($\mu\text{s}$) | $\alpha_2$ | $t_2$ (ms) | $\alpha_3$ | $t_3$ (ms) | $\alpha_4$ | $t_4$ (s) | $\alpha_5$ | $t_5$ (s) |
| --- | --- | --- | --- | --- | --- | --- | --- | --- | --- | --- |
| +1 (iCy3)<sub>2</sub> dimer, [NaCl] = 300 mM, [MgCl<sub>2</sub>] = 6 mM | 0.90 | 250 | 0.24 | 2.2 | 0.18 | 18 | 0.14 | 0.15 | 0.20 | 2.0 |
| +1 (iCy3)<sub>2</sub> dimer, [NaCl] = 100 mM, [MgCl<sub>2</sub>] = 6 mM | 0.8 | 300 | 0.18 | 3.7 | 0.20 | 170 | 0.30 | 3.1 | NA | NA |
| +1 (iCy3)<sub>2</sub> dimer, [NaCl] = 20 mM, [MgCl<sub>2</sub>] = 6 mM | 1.0 | 270 | 0.21 | 2.8 | 0.13 | 90 | 0.23 | 1.5 | NA | NA |
| +1 (iCy3)<sub>2</sub> dimer, [NaCl] = 20 mM, [MgCl<sub>2</sub>] = 0 mM | 1.0 | 290 | 0.26 | 3.7 | 0.19 | 82 | 0.12 | 1.1 | NA | NA |
| -1 (iCy3)<sub>2</sub> dimer, [NaCl] = 100 mM, [MgCl<sub>2</sub>] = 6 mM | 0.62 | 600 | 0.27 | 2.2 | 0.13 | 53 | 0.14 | 1.4 | NA | NA |
| -2 (iCy3)<sub>2</sub> dimer, [NaCl] = 300 mM, [MgCl<sub>2</sub>] = 6 mM | 1.0 | 360 | 0.13 | 5.1 | 0.12 | 200 | 0.17 | 2.5 | NA | NA |
| -2 (iCy3)<sub>2</sub> dimer, [NaCl] = 100 mM, [MgCl<sub>2</sub>] = 6 mM | 1.0 | 390 | 0.19 | 3.0 | 0.12 | 93 | 0.08 | 2.0 | NA | NA |
| -2 (iCy3)<sub>2</sub> dimer, [NaCl] = 20 mM, [MgCl<sub>2</sub>] = 6 mM | 1.0 | 310 | 0.18 | 5.9 | 0.13 | 100 | 0.21 | 1.6 | NA | NA |
| -2 (iCy3)<sub>2</sub> dimer, [NaCl] = 20 mM, [MgCl<sub>2</sub>] = 0 mM | 1.0 | 360 | 0.20 | 2.9 | 0.10 | 22.0 | 0.07 | 1.2 | 0.12 | 4.6 |

**Figure 9.**
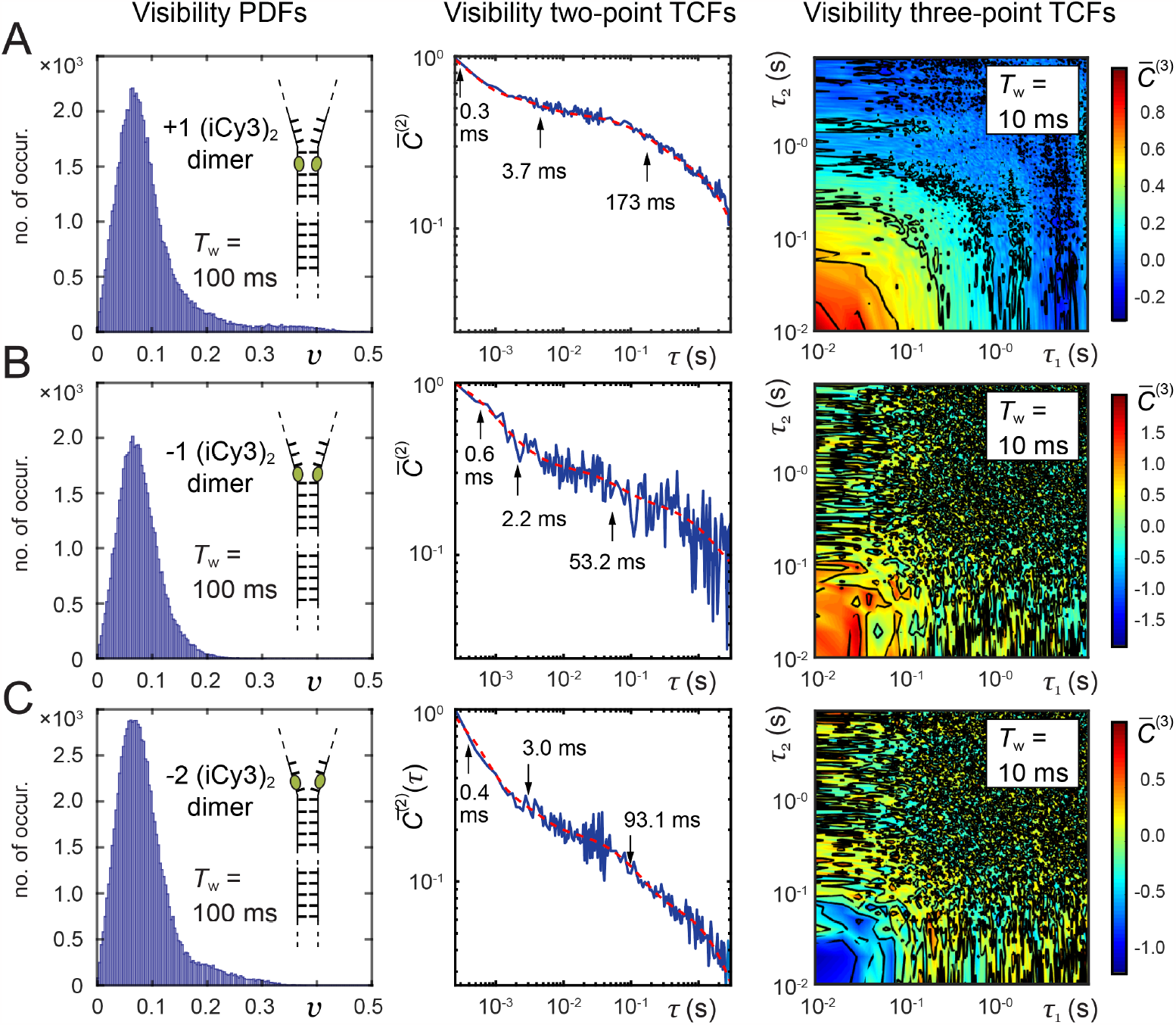
Probability distribution functions (PDFs, left column), two-point time-correlation functions (TCFs, middle column) and three-point TCFs (left column) of the PS-SMF visibility for the (panel ***A***) +1, (panel ***B***) -1 and (panel ***C***) -2 (iCy3)_2_ dimer-labeled ss-dsDNA fork constructs. The integration windows used for the PDFs and three-point TCFs are indicated in the insets. The two-point TCFs were stitched together over the range 250 μs – 25 ms using *T*_*w*_ = 250 μs, and from 25 ms – 2.5 s using *T*_*w*_ = 10 ms. The two-point TCFs are shown overlaid with multi-exponential fits (dashed red curves) whose parameters are listed in Table 3. The three-point TCFs are plotted as two-dimensional contour diagrams. Experiments were performed at room temperature (23 °C) and physiological buffer salt conditions ([NaCl] = 100 mM, [MgCl_2_] = 6 mM).

In the second and third columns of Fig. 9, respectively, we present the two-point and three-point TCFs, which characterize the conformational dynamics of the (iCy3)_2_ dimer-labeled ss-dsDNA constructs as a function of probe labeling position. The two-point TCFs are shown overlaid with multi-exponential fits (dashed red curves) whose parameters are listed in Table 3. For each of the (iCy3)_2_ dimer-labeled ss-dsDNA constructs, the four decay components are well-separated in time: *t*_1_ ∼0.3 – 0.6 ms, *t*_*2*_ ∼2 – 4 ms, *t*_3_ ∼50 -200 ms and *t*_4_ ∼1 – 2 s. As mentioned previously, the seconds-long decay, *t*_4_, is due to mechanical room vibrations. We thus find that there are three relevant, well-separated decay components present in all the (iCy3)_2_ dimer-labeled ss-dsDNA constructs studied under physiological salt conditions, as well as at the higher and lower salt concentration conditions that we also examined in this study (see Table 3).

From the two-point TCFs of Fig. 9, we see that the + 1 construct exhibits the slowest overall decay (Fig. 9*A*). As the position of the (iCy3)_2_ dimer is varied across the ss-dsDNA junction from +1 to -1 (see Fig. 9*B*), and again from -1 to -2 (Fig. 9*C*), the relaxation dynamics become faster. These observations are consistent with the notion that the conformational motions of the (iCy3)_2_ dimer probe depend on the stabilities and dynamics of the DNA bases and sugar-phosphate backbones immediately adjacent to the probe. The Watson-Crick (WC) base pairs within the duplex side of the ss-dsDNA junction are largely stacked, while of course the stacking interactions between bases on the ssDNA side of the junction are much weaker due to the lack of complementary base pairing. We note that the three-point TCFs, 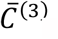, exhibit positive amplitude for the +1 and -1 constructs, and negative amplitude for the -2 construct at short times (10 ms < τ_1_, τ_*2*_ < 100 ms). These observations suggest that the dominant kinetic pathways that govern conformational transitions for the +1 and -1 constructs, which undergo exchange dynamics relatively slowly, are distinct from the pathways that govern the transitions of the -2 construct, which undergoes relatively fast exchange dynamics.

### 3.3 Salt concentration-dependent breathing of +1 and -2 (iCy3)_2_ dimer-labeled ss-dsDNA fork constructs

We next consider the effects of varying buffer salt concentration on the equilibrium properties and dynamics of the (iCy3)_2_ dimer-labeled ss-dsDNA fork constructs. We examined the effects of varying salt concentration on the slowest (+1 position) and the fastest (-2 position) of the ss-dsDNA constructs (see Fig. 10 and Fig. 11, respectively). In the case of the +1 construct the dimer probe is positioned within the dsDNA region immediately adjacent to the ss-dsDNA fork junction, while the -2 construct has the dimer probe positioned within the ssDNA region immediately adjacent to the junction.

**Figure 10.**
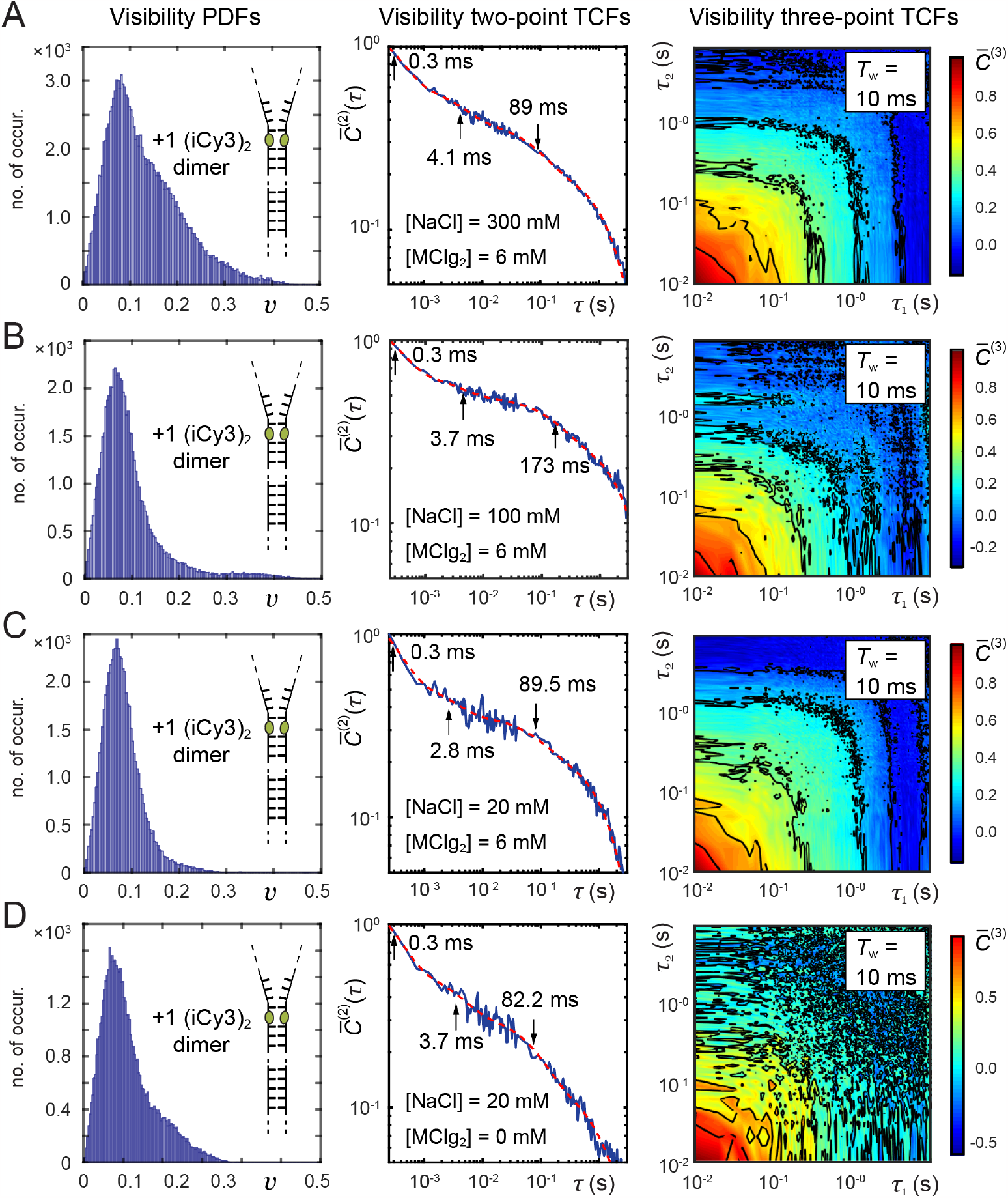
Salt concentration-dependent PDFs (left column), two-point TCFs (middle column) and three-point TCFs (right column) of the signal visibility for the +1 (iCy3)_2_ dimer-labeled ss-dsDNA construct. Experiments were performed at room temperature (23°C) and using (panel ***A***) [NaCl] = 300 mM, [MgCl_2_] = 6 mM; (panel ***B***) [NaCl] = 100 mM, [MgCl_2_] = 6 mM; (panel ***C***) [NaCl] = 20 mM, [MgCl_2_] = 6 mM; and (panel ***D***) [NaCl] = 20 mM, [MgCl_2_] = 0 mM.

**Figure 11.**
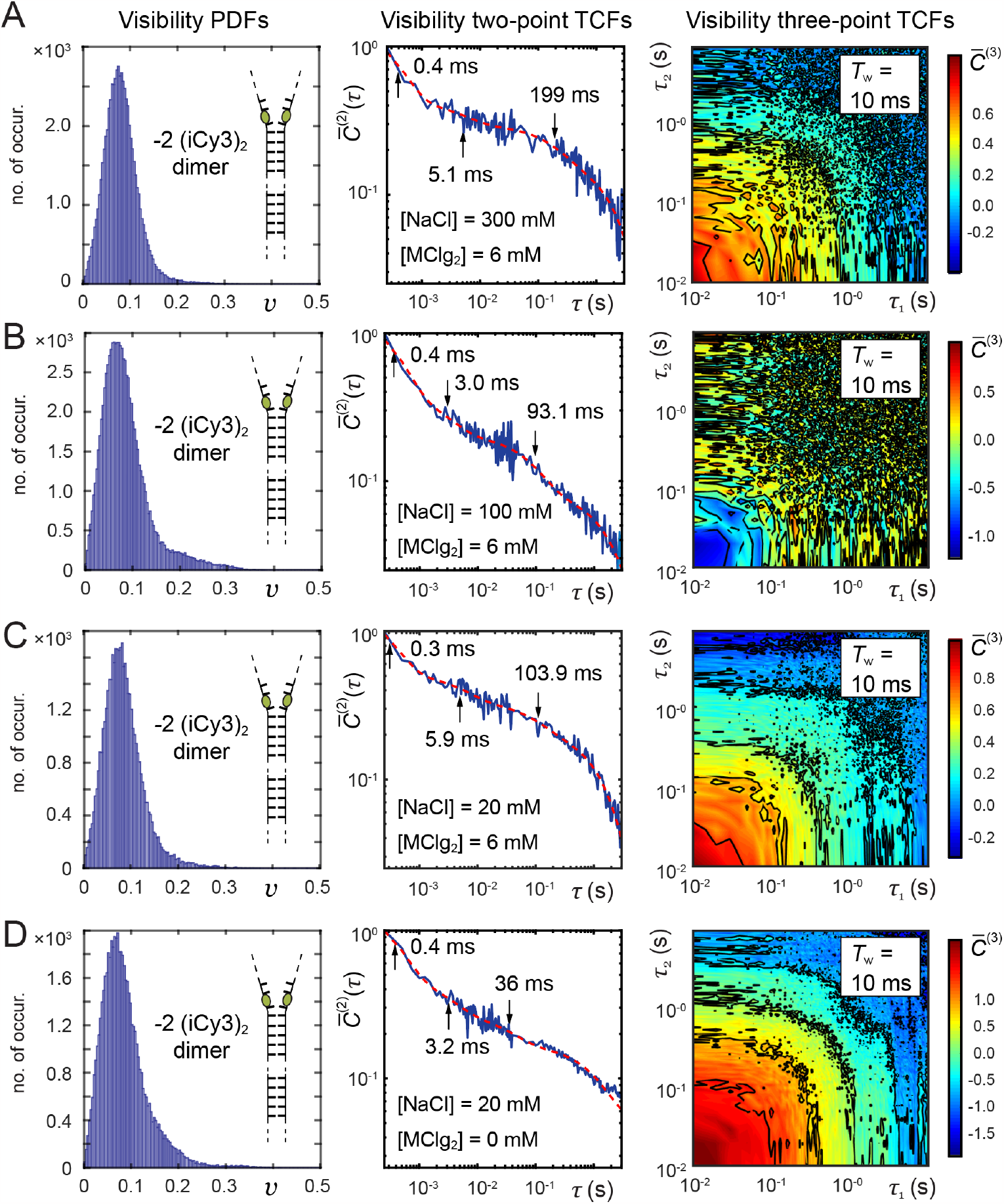
Salt concentration-dependent PDFs (left column), two-point TCFs (middle column) and three-point TCFs (right column) of the signal visibility for the -2 (iCy3)_2_ dimer-labeled ss-dsDNA construct. Experiments were performed at room temperature (23°C) and using (panel ***A***) [NaCl] = 300 mM, [MgCl_2_] = 6 mM; (panel ***B***) [NaCl] = 100 mM, [MgCl_2_] = 6 mM; (panel ***C***) [NaCl] = 20 mM, [MgCl_2_] = 6 mM; and (panel ***D***) [NaCl] = 20 mM, [MgCl_2_] = 0 mM.

It is interesting to compare the differing effects of salt concentration on our DNA constructs as reflected in the structures of the (iCy3)_2_ dimer probes as a function of probe position at and near the ss-dsDNA forks. Based on our previous ensemble studies carried out at physiological salt conditions ([NaCl] = 100 mM, [MgCl_2_] = 6 mM) the DNA duplex is thought to adopt primarily the Watson-Crick right-handed conformation, although thermally excited conformational macrostates may be populated transiently. The marginal stability of the DNA duplex results from a near balance between opposing thermodynamic forces (i.e., enthalpy-entropy compensation) [21, 32]. For example, we previously concluded [11] that the forces that favor base-stacking are due primarily to the positive solvent entropy changes that accompany the expulsion of the water molecules that are otherwise confined between flat base surfaces (i.e., the depletion effect) [33, 34].

The forces that oppose base stacking are enthalpic. These reflect primarily the strain on the sugar-phosphate backbones due to the electrostatic repulsion between negatively charged phosphate backbone groups, which are approximately 70*w* screened by the diffuse ‘ion cloud’ of monovalent sodium ions within the condensation layer immediately surrounding the negatively charged DNA-water interface under physiological salt concentrations. Decreasing the salt concentration from physiological levels destabilizes the DNA duplex by reducing the electrostatic screening between negatively charged phosphate groups [22-28]. On the other hand, increasing salt concentration above physiological conditions can also destabilize the DNA duplex. The origins of duplex destabilization at moderately elevated salt concentrations (> 100 mM NaCl) are not fully understood, although this may be partly due to a reduction of the water entropy associated with the formation of structured solvation shells around increasing concentrations of ions and counterions in bulk solution. At salt concentrations much higher than physiological (∼ 1M), the duplex may be either stabilized or destabilized through specific Hoffmeister ion effects on macromolecular surfaces in aqueous solutions [35, 36]. Few details are currently available about how salt concentration affects the equilibrium distribution of conformational macrostates at and near ss-dsDNA fork junctions, or the transition barriers that mediate their interconversion.

In Fig. 10, we present the results of our salt concentration-dependent studies for the +1 (iCy3)_2_ dimer-labeled ss-dsDNA construct. These results are organized, from top row to bottom, in order of decreasing salt concentration. We note that for all the salt concentrations that we studied, the PDFs exhibit a pattern of low and high visibility features, and most two-point and three-point TCFs exhibit three well-separated decay components, like those discussed in previous sections. An exception is the TCF corresponding to the highest salt condition shown in Fig. 10*A*, which exhibits four well-separated decay components (see Table 3). As described in previous ensemble work [11, 12], the presence of three well-separated decay components suggests that the simplest kinetic network model that can be used to simulate these systems involves four macrostates in thermal equilibrium.

At the highest salt concentrations that we studied ([NaCl] = 300 mM, [MgCl_2_] = 6 mM, Fig. 10*A*), the PDF of the +1 (iCy3)_2_ dimer-labeled ss-dsDNA construct exhibits increased population of ‘high visibility’ macrostates within the 0.15 ≲ *v* ≲ 0.45 range in comparison to the PDFs measured at physiological salt conditions (see Fig. 10*B*). Moreover, the TCFs at high salt concentration decay significantly faster, suggesting that the heights of the transition barriers for state-to-state interconversion are reduced under these conditions. When the salt concentration is decreased (relative to physiological conditions) to [NaCl] = 20 mM, [MgCl_2_] = 6 mM (Fig. 10*C*), the PDF exhibits a slight narrowing of the distribution of ‘low visibility’ macrostates within the 0 ≲ *v* ≲ 0.15 range, and a reduced population of ‘high visibility’ macrostates within the 0.25 ≲ *v* ≲ 0.45 range. Furthermore, the TCFs exhibit faster dynamics at these low salt concentrations, suggesting that the most heavily weighted macrostates exhibit shorter lifetimes in comparison to physiological conditions.

Finally, elimination of divalent magnesium ions at the lowest sodium concentration ([NaCl] = 20 mM, [MgCl_2_] = 0 mM, Fig. 9*D*) leads to the reappearance in the PDF of ‘high visibility’ macrostates in the range 0.1 ≲ *v* ≲ 0.3, albeit with a Boltzmann-weighted distribution quite different than observed at physiological conditions. The dynamics exhibited by the TCFs are fast, suggesting that the transition barriers of the most heavily weighted macrostates are reduced relative to physiological salt conditions. In summary, the effects of raising and lowering salt concentration from physiological conditions for the +1 (iCy3)_2_ dimer-labeled ss-dsDNA construct are to shift the equilibrium balance of ‘high’ and ‘low visibility’ conformational macrostates to favor ‘high visibility’ states and to lower the free energy of activation for state-to-state interconversion.

We next examined the effects of varying salt concentration on the -2 (iCy3)_2_ dimer-labeled ss-dsDNA construct (see Fig. 11). The TCF of the -2 DNA construct at physiological salt concentrations exhibits three well-separated decay components, which are significantly faster than for the +1 construct. Moreover, the PDF of the -2 construct at physiological salt concentrations exhibits significant population of ‘high visibility’ states within the 0.15 ≲ *v* ≲ 0.35 range, indicating that the bases and sugar-phosphate backbones immediately adjacent to the dimer probes at the -2 position adopt a different distribution of conformational macrostates than those of the +1 (iCy3)_2_ dimer-labeled ss-dsDNA construct. The relatively fast dynamics of the -2 construct indicates that the transition barriers of the most heavily weighted macrostates are relatively low compared to the +1 construct.

Interestingly, the effects of increasing and decreasing salt concentration on the -2 dimer-labeled DNA construct relative to physiological conditions are opposite to those we observed for the +1 construct. At elevated monovalent salt concentration ([NaCl] = 300 mM, [MgCl_2_] = 6 mM, Fig. 11*A*), the PDF exhibits a narrowing of the ‘low visibility’ states within the 0 ≲ *v* ≲ 0.15 range and reduced population of ‘high visibility’ states (Fig. 11*A*). Moreover, the relaxation dynamics are significantly slowed, suggesting that the activation barriers for state-to-state interconversion are elevated. A similar effect occurs for reduced monovalent salt concentration ([NaCl] = 20 mM, [MgCl_2_] = 6 mM, Fig. 11*C*). Elimination of divalent magnesium ions at the reduced sodium ion concentration ([NaCl] = 20 mM, [MgCl_2_] = 0 mM, Fig. 11*D*) leads to a slight broadening of the PDF and faster relaxation dynamics, like those observed at physiological salt conditions. In addition, at this low salt concentration the TCFs exhibit four well-separated decay components (see Table 3). We note that the primary negative amplitude feature of the three-point TCF, seen at physiological salt concentrations, becomes positive at elevated and reduced salt concentrations under conditions in which the dynamics have slowed. These observations indicate that the effects of varying salt concentration relative to physiological conditions for the -2 (iCy3)_2_ dimer-labeled ss-dsDNA construct are to shift the distribution of conformational macrostates to favor ‘low visibility’ conformational macrostates and to raise the free energy of activation for state-to-state interconversion.

## 4. Conclusions

In this work, we have introduced a novel experimental method, called polarization-sweep single-molecule fluorescence (PS-SMF) microscopy, to study site-specific DNA ‘breathing’ fluctuations at and near ss-dsDNA fork junctions. PS-SMF uses exciton-coupled (iCy3)_2_ dimer-labeled ss-dsDNA constructs to directly monitor the fluctuations of the dimer probes, which depend sensitively on the local conformations of the DNA bases and sugar-phosphate backbones immediately adjacent to the dimer probes at specific site positions.

Our results (summarized in Fig. 9 – Fig. 11) indicate that the bases and sugar-phosphate backbones sensed by the (iCy3)_2_ dimer probes can adopt four quasi-stable local conformations, whose relative stabilities and transition state barriers depend on probe labeling position relative to the ss-dsDNA fork junction. Under physiological buffer salt conditions ([NaCl] = 100 mM, [MgCl_2_] = 6 mM), the +1 (iCy3)_2_ dimer-labeled ss-dsDNA construct exhibits a relatively broad distribution of local base and backbone conformations within the duplex region of the ss-dsDNA junction (Fig. 9). The conformational macrostates at this position undergo the slowest dynamics of all the systems that we investigated, indicating that the most thermodynamically stable states within the duplex side of the fork junction are also mechanically stable with relatively long population lifetimes. In comparison, the -2 dimer-labeled ss-dsDNA construct exhibits significantly faster dynamics and a distinctly different distribution of conformational macrostates, indicating that the thermodynamically favored conformations of bases and sugar-phosphate backbones sensed by the dimer probes within the ssDNA region of the fork junction are mechanically unstable and distinct from the duplex.

The energetics of the (iCy3)_2_ dimer-labeled ss-dsDNA constructs can be systematically varied by increasing or decreasing salt concentrations relative to physiological conditions. The position-dependent distribution of conformational macrostates is affected by salt concentration in complementary ways. For the +1 construct, increasing or decreasing salt concentration relative to physiological conditions shifts the equilibrium distribution to favor macrostates that are mechanically unstable (with relatively low transition barriers, see Fig. 10). For the -2 construct, varying salt concentration shifts the equilibrium distribution of macrostates to favor those that are mechanically stable (with relatively high transition barriers, see Fig. 11).

In Fig. 12, we illustrate a hypothetical mechanism to account for these observations. In the case of the +1 dimer-labeled ss-dsDNA construct, the local base and backbone conformations adjacent to the dimer probes are within the DNA duplex region of the fork junction (Fig. 12*A*). At physiological salt conditions, the majority of conformational macrostates sensed by the dimer probe are dominated by WC base stacking, and these conformations are mechanically stable. Increasing or decreasing salt concentration leads to disruption of the local WC conformations within the duplex region and the reduction of mechanical stability (Fig. 12*B*). In the case of the -2 dimer-labeled ss-dsDNA construct, the local base and backbone conformations adjacent to the dimer probes are within the ssDNA region (Fig. 12*C*). At physiological salt conditions, the majority of conformational macrostates sensed by the dimer in the ssDNA region are dominated by unstacked base conformations, which are distinct from the duplex region and mechanically unstable. Further evidence for the presence of unstacked and dynamically labile base conformations within the ssDNA region of oligo(dT)_15_ tails of ss-dsDNA fork constructs was determined in microsecond-resolved single-molecule FRET experiments [11] and corroborated by small-angle x-ray scattering experiments [37, 38]. At elevated or reduced salt concentration, the distribution of conformational macrostates sensed by the dimer probes within the ssDNA region of the fork junction is shifted to mechanically stable base-stacked conformations (Fig. 12*D*). The above hypothetical mechanism is consistent with the findings of previous ensemble studies of (iCy3)_2_ dimer labeled ss-dsDNA fork constructs [6], and are further interpreted using kinetic network modeling approaches in a subsequent paper [10].

**Figure 12.**
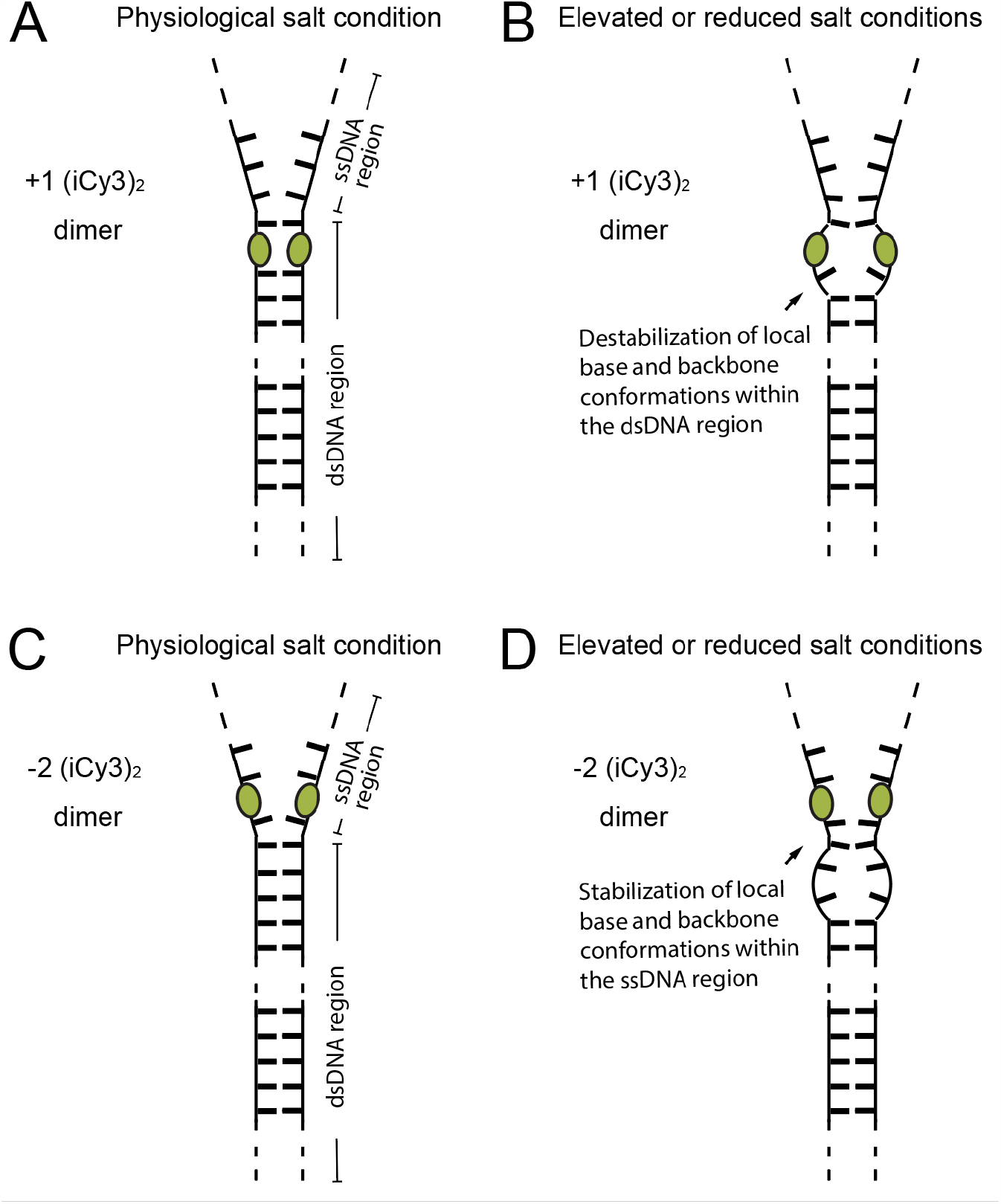
Schematic diagram illustrating our hypothesized mechanism of salt-induced instability of (iCy3)_2_ dimer-labeled ss-dsDNA fork constructs labeled at (panel ***A*** and ***B***) the +1 position and (panel ***C*** and ***D***) the -2 position. Left column diagrams (panels ***A*** and ***C***) represent the situation at physiological salt conditions, and diagrams in the right column (panels ***B*** and ***D***) represent our hypothesized mechanisms at elevated or reduced salt conditions.

PS-SMF data, such as those presented in the current work, can be further analyzed using a kinetic network model [11-13] to provide quantitative information about the relative stabilities of the various macrostates and their free energies of activation. The structural and kinetic properties of the ss-dsDNA fork junction revealed by these experiments can provide mechanistic insights about how replication and repair proteins that assemble at these sites carry out their biological functions.

The PS-SMF method monitors the signal visibility, *v*, from exciton-coupled (iCy3)_2_ dimer probes that are rigidly inserted within the sugar-phosphate backbones of the ss-dsDNA fork construct. Unlike single-molecule optical experiments that solely monitor fluorescence intensity, the visibility observable is a reduced quantity that is directly related to the alignment of the monomeric subunits of the dimer probe. Therefore, PS-SMF is a useful means to measure local structure at the single-molecule level on the scale of a few Angstroms, which cannot be otherwise studied using other single-molecule optical approaches. The PS-SMF experiments presented here focus on local DNA conformations at and near model ss-dsDNA fork junctions, which are relevant to the non-base-sequence specific assembly of replication and repair proteins at these sites. However, the PS-SMF approach could be applied advantageously to study the local conformations and conformational fluctuations relevant to base-sequence specific recognition of proteins that function to regulate genes, in addition to the thermodynamic and kinetic effects of specific covalent modifications of DNA bases (e.g., see [39, 40]) that can lead to epigenetic (non-base-pairing structural) effects at the DNA level.

## Acknowledgements

The authors are grateful to their laboratory colleagues for many helpful discussions. This work was supported by grants from the National Institutes of Health General Medical Sciences (Grant No. GM-15792 to A.H.M. and P.H.v.H.), and the National Science Foundation RAISE-TAQS Program (Grant No. PHY-1839216 to A.H.M.). P.H.v.H. is an American Cancer Society Research Professor of Chemistry.

## Appendices Appendix A: Polarization-sweep single-molecule fluorescence (PS-SMF) interferometer

In Fig. S1, we show a schematic of the laser interferometer and the integrated detection electronics used for our PS-SMF experiments. A half-wave plate is used to rotate the plane polarization vector of a continuous wave (cw) 532 nm laser to 45° from vertical. The laser is directed through a polarizing beam-splitter (PBS) to produce balanced vertical and horizontal plane-polarized beams, which traverse separately the two paths of a Mach-Zehnder Interferometer (MZI) before they are recombined at the PBS exit port. Two acousto-optic Bragg cells (AOBCs) are each placed within the paths of the MZI. Each AOBC is driven continuously at a fixed radio frequency so that a relative phase ‘sweep’ is imparted to the MZI output beam: φ = Ω*t* with Ω = 1 MHz. The output beam passes through a quarter-wave plate, which is rotated to 45° from vertical, before it is directed to the TIRF microscope to excite the single-molecule sample (see Fig. 2*A* of the main text). A pick-off mirror is used to direct a minor component of the beam through a polarizer and detected using an avalanche photodiode (APD) to create a 1 MHz analog reference signal. The reference is utilized in a negative feedback loop to actively minimize the path difference of the MZI using a piezo-controlled mirror, as implemented in a previous experiment [17].

**Figure A1.**
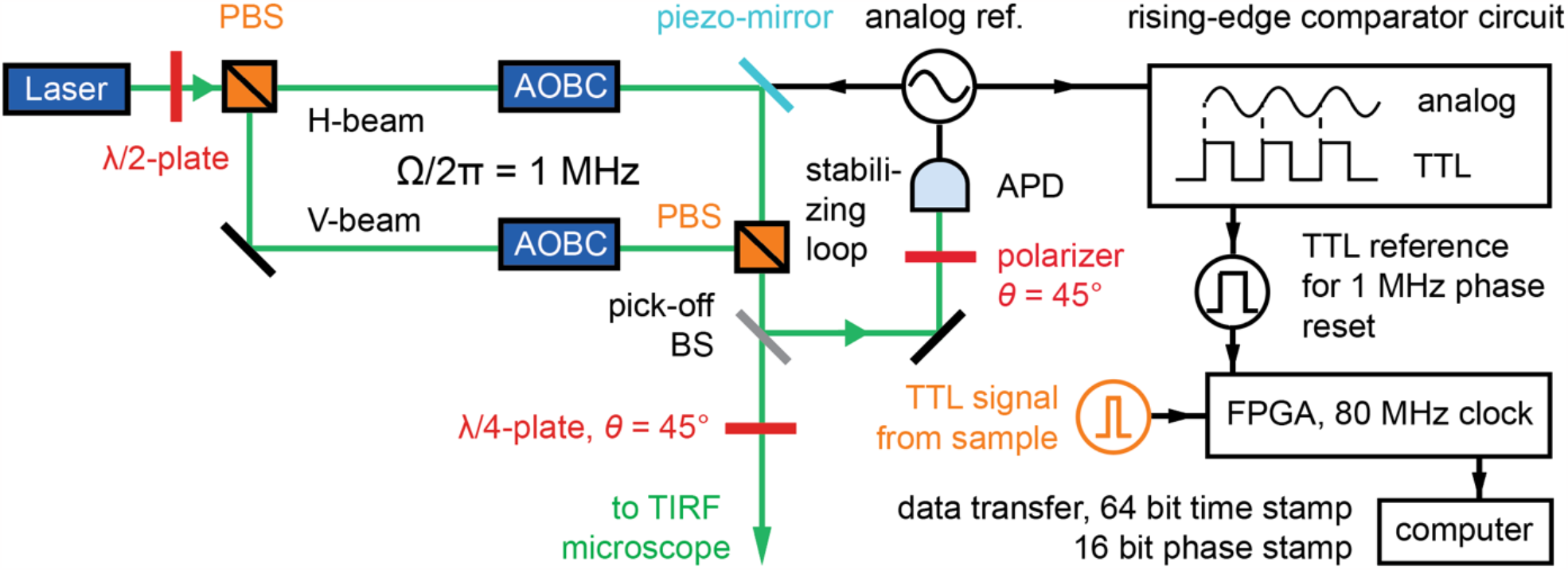
Schematic of the interferometer and integrated phase-tagged photon counting (PTPC) electronics [8] used to perform PS-SMF experiments. PBS: polarizing beam-splitter; AOBC: acousto-optic Bragg cell; APD: avalanche photodiode; FPGA: field-programmable gate array.

The digital electronics illustrated in Fig. A1 are used to implement the phase-tagged photon counting (PTPC) method described by [8]. In this approach, we use a field-programmable gate array (FPGA), which is a manually reconfigurable integrated circuit that contains hardware-enabled signal-processing algorithms. We used the FPGA to discretize the phase of a given modulation cycle into a set of *m* ‘phase bins,’ which are numbered and incrementally advanced using an 80 MHz digital counter. The 1 MHz analog reference waveform is first converted into a ‘logical square wave’ (i.e., a periodic TTL waveform) using a custom-built rising-edge comparator circuit. The logical square wave is then used to trigger an 80 MHz phase counter (16-bit width), and to reset the counter at the 1 MHz phase-sweep frequency. The counter reset automatically synchronizes the phase-sweep cycle in the presence of external room vibrations that introduce noise to the MZI reference phase. The average number of phase bins during a modulation cycle is *m* = 80 MHz / 1 MHz = 80 bins, and the phase bin interval is Δφ = 360° / 80 = 4.5° bin^-1^. Thus, during each phase-sweep cycle the counter advances the phase bin value φ_j_ = *j*Δφ (with *j* = 0, 1, …, 79) at the 1 MHz clock speed. We also used a second 1 MHz counter (with 64bit width) to assign a time bin value *t*_k_ = *k*Δ*t* (with Δ*t* = 1 μs and *k* = 0, 1, …) to each detection event. Thus, each individual detection event is assigned a 16-bit phase bin value and a 64-bit time bin value, and these data are streamed to computer memory in real time.

## Appendix B Theoretical description of the rotating plane-polarized optical field

In this section, we derive Eq. (1) of the main text. We decompose the polarization state of the electric field in the basis of horizontal (*H*) and vertical (*V*) plane polarization components. Before entering the MZI (see Fig. A1), the plane polarization of the laser is rotated to 45° from vertical using a half-wave plate so that the initial electric field may be written in the 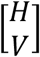tbasis:

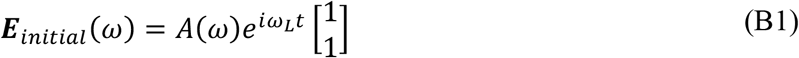

In Eq. (B1), *A*(ω) is a narrow spectral envelope and ω_G_ is the laser center frequency (= 2π*c*/λ_*L*_, where λ_*L*_ = 532 nm). At the entrance port of the MZI, the PBS separates the *H* and *V* polarization components into separate paths. Within each MZI path, an AOBC imparts a time-varying phase shift to its respective beam according to φ_H_ = Ω_H_*t* and φ_*V*_ = Ω_*V*_*t*. Thus, the electric field components within the *H* and *V* paths can be written:

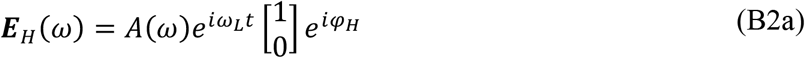

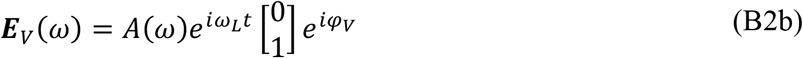

At the exit port of the MZI, the *H* and *V* beams are recombined to produce the total field

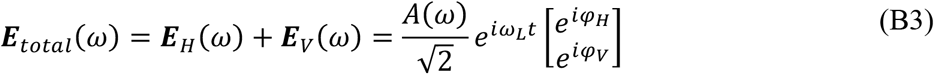

Equation (B3) can be simplified by defining the relative phase shift φ = φ_V_ − φ_H_ = Ω_VH_*t* between the *H* and *V* paths (with Ω_VH_ = Ω = 1 MHz), and by multiplying by the overall phase shift *e*−*i*(ϕ*V*_+_ϕH)/*2*:

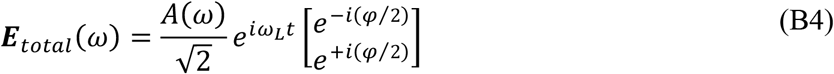

The beam is next directed through a quarter-wave plate that is rotated 45° from vertical. The ‘Jones matrix’ *M* for the quarter-wave plate is given by

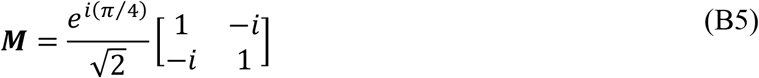

Thus, after passing through the quarter-wave plate, the final electric field can be written compactly as:

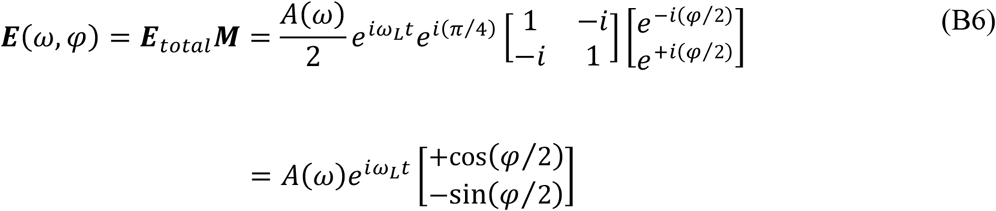

The second equality of Eq. (B6) is the electric field incident on the sample, which is equivalent to Eq. (1) of the main text.

## Appendix C Derivation of the polarized signal intensity for PS-SMF experiments

Here we derive Eq. (4) of the main text. The electric field vector, *E*(ω, φ), is given by Eq. (B6) and the symmetric (+) and anti-symmetric (–) electric dipole transition moments (EDTMs) of the exciton coupled dimer, μ_±_(ω), are given by

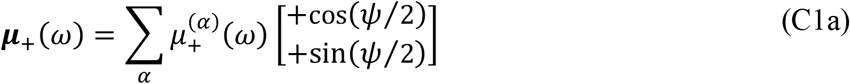

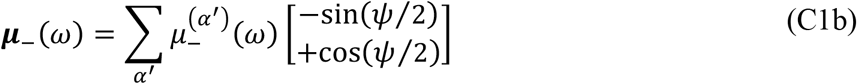

In Eqs. (C1a) and (C1b), the angle ψ/2 specifies the orientation of the dimer in the *x*-*y* plane, 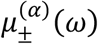 are the spectrally-dependent magnitudes of the EDTMs, which depend on the dimer conformation, and the index α enumerates the singly-excited dimer states in order of increasing energy [4-6]. The μ_±_ symmetric and anti-symmetric EDTMs (μ_+_ and μ___, respectively) are depicted graphically in Fig. 2*C* of the main text. At any instant, the ‘polarized’ single-molecule fluorescence intensity is given by the spectral overlap and square modulus of the vector projections between the laser electric field and the total EDTM.

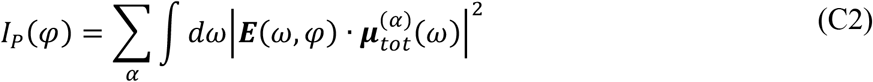

Equation (C2) can be written as the sum of symmetric (+) and anti-symmetric (–) exciton contributions by substituting μ_*tot*_ = μ_+_ + μ___:

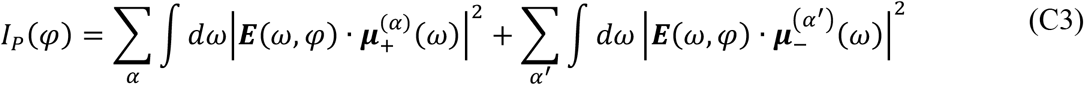

We identify the first term on the right-hand side of Eq. (C3) as the symmetric exciton contribution, *I*_+_(φ), and the second term as the anti-symmetric exciton contribution, *I*___(φ). The separation given by Eq. (C3) is a consequence of μ_+_ and μ___ having orthogonal orientations, which additionally leads to the two signal contributions having a relative phase of 180°.

We first calculate the signal contribution from the symmetric excitons. Substitution of Eq. (B6) and Eq. (C1a) into the first term on the right-hand side of Eq. (C3) leads to:

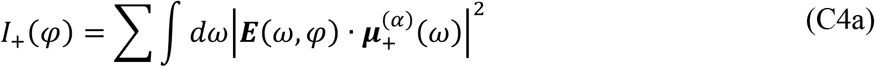

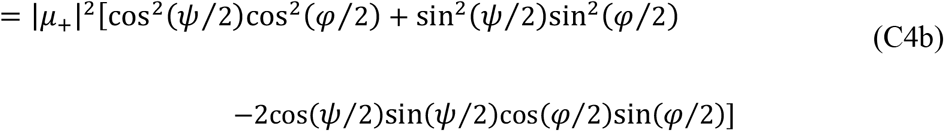

where we have defined the spectral overlap function 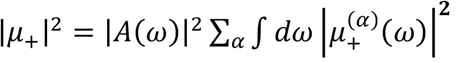.

Equation (C4b) can be simplified using double angle formulas to yield

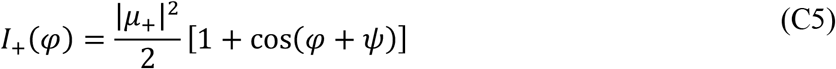

Following a similar procedure for the anti-symmetric exciton contribution to the polarized signal [by substitution of Eq. (B6) and Eq. (C1b) into the second term on the right-hand side of Eq. (C3)] leads to:

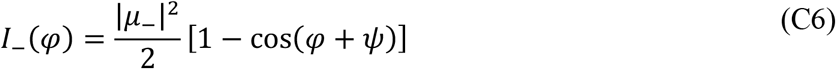

where 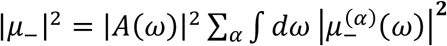. Combining Eqs. (C5) and (C6) gives the expression for the polarized single-molecule fluorescence intensity:

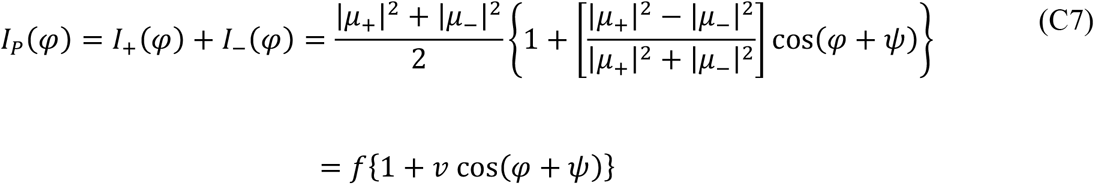

In Eq. (C7), 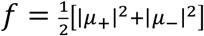 is the mean signal rate and *v* = [|μ_+_|^*2*^ − |μ___|^*2*^]/[|μ_+_|^*2*^ + |μ___|^*2*^] is the signal visibility.

---

**Figure S1.**
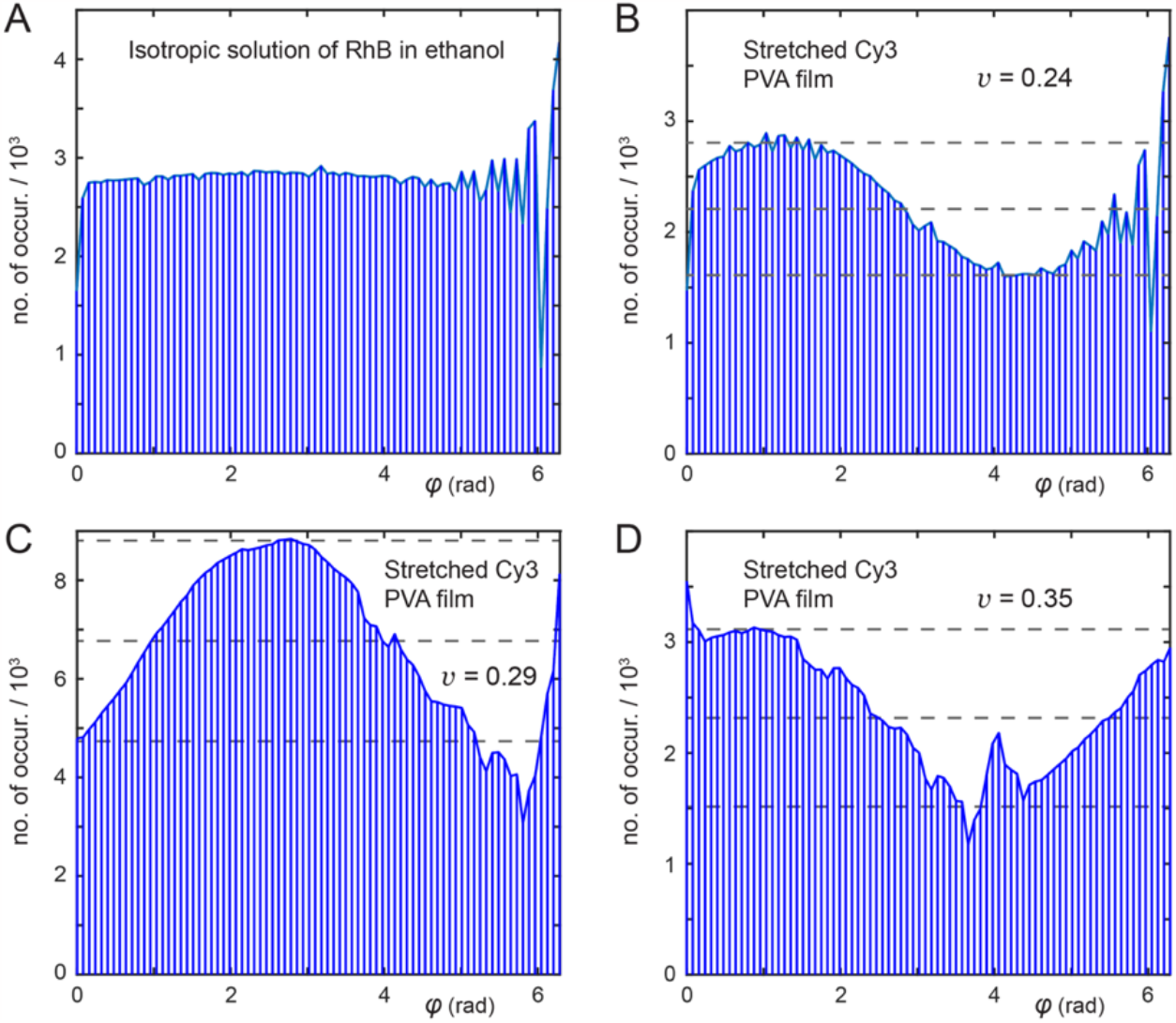
Control measurements of phase histograms from (***A***) an isotropic solution of Rhodamine B in ethanol, and (***B*** – ***D***) an anisotropically-stretched film of Cy3 supported in poly vinyl alcohol (PVA). For each of the panels, ***B*** – ***D***, the uniaxial stretch direction and spot location of the PVA film was varied. Histograms were constructed from an average of 10 separate time series, each of ∼45 s duration. For visualization, the histograms were constructed by re-binning the native 16,000 bin histograms into 79 bins, so that the resolution shown is course-grained to ∼ 0.08 rad bin^-1^. The horizontal dashed lines indicate the maximum (top), mean (middle) and minimum (bottom) number of counts per bin. The signal was detected using the phase-tagged photon counting (PTPC) method [1].

**Figure S2.**
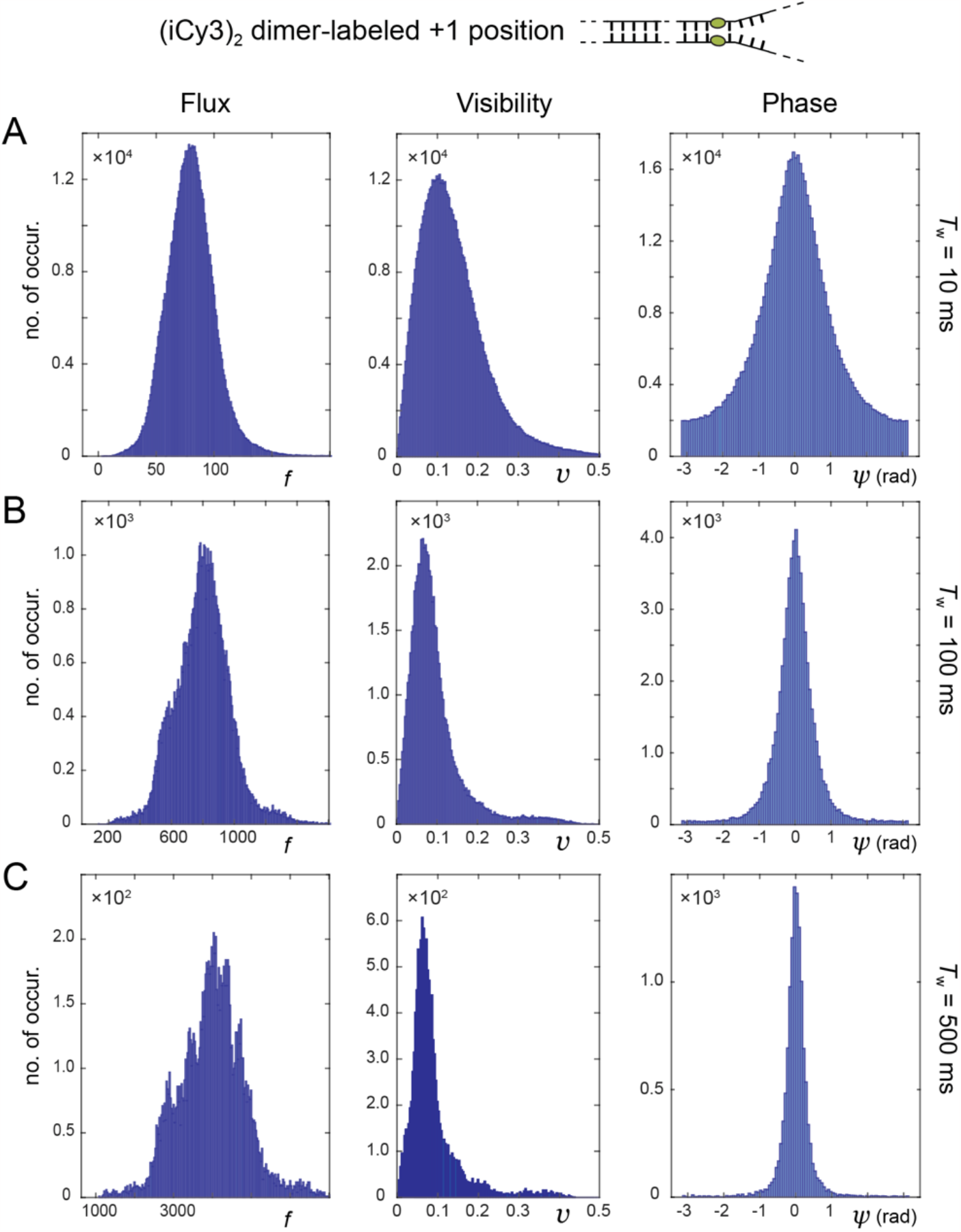
PS-SMF probability distribution functions (PDFs) for the +1 (iCy3)_2_ dimer-labeled ss-dsDNA fork construct. Histograms were constructed from the raw photon data streams (see, e.g., Fig. 3 of the main text) for the signal flux (left column), signal visibility (middle column) and signal phase (right column). The mean signal flux 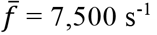. Comparisons are shown varying the integration window (***A***) *T*_w_ = 10 ms, (***B***) *T*_w_ = 100 ms and (***C***) *T*w = 500 ms. The corresponding 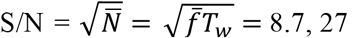 and 61, respectively.

**Figure S3.**
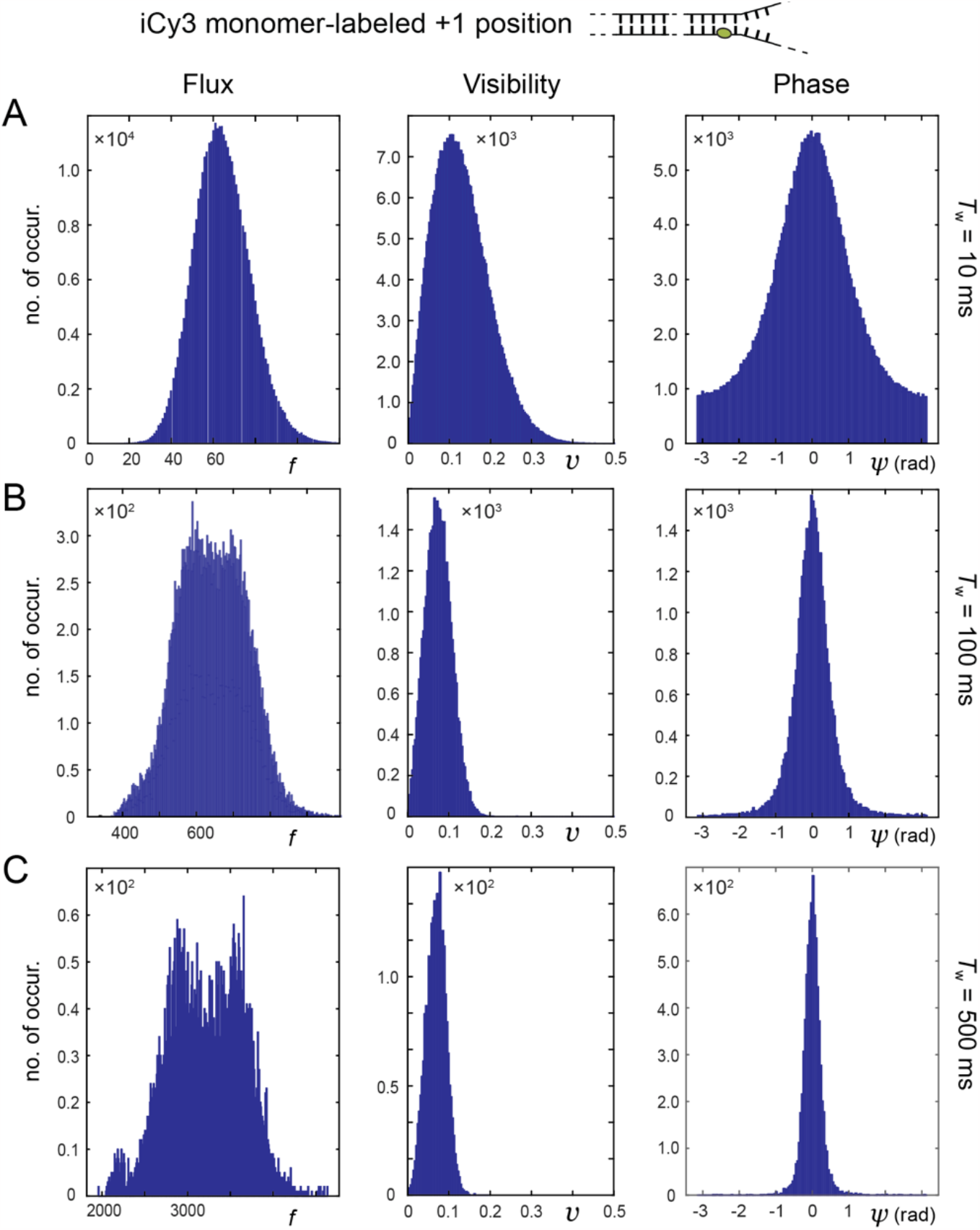
PS-SMF probability distribution functions (PDFs) for the +1 iCy3 monomer-labeled ss-dsDNA fork construct. Panels are the same as described in Fig. S2.

**Figure S4.**
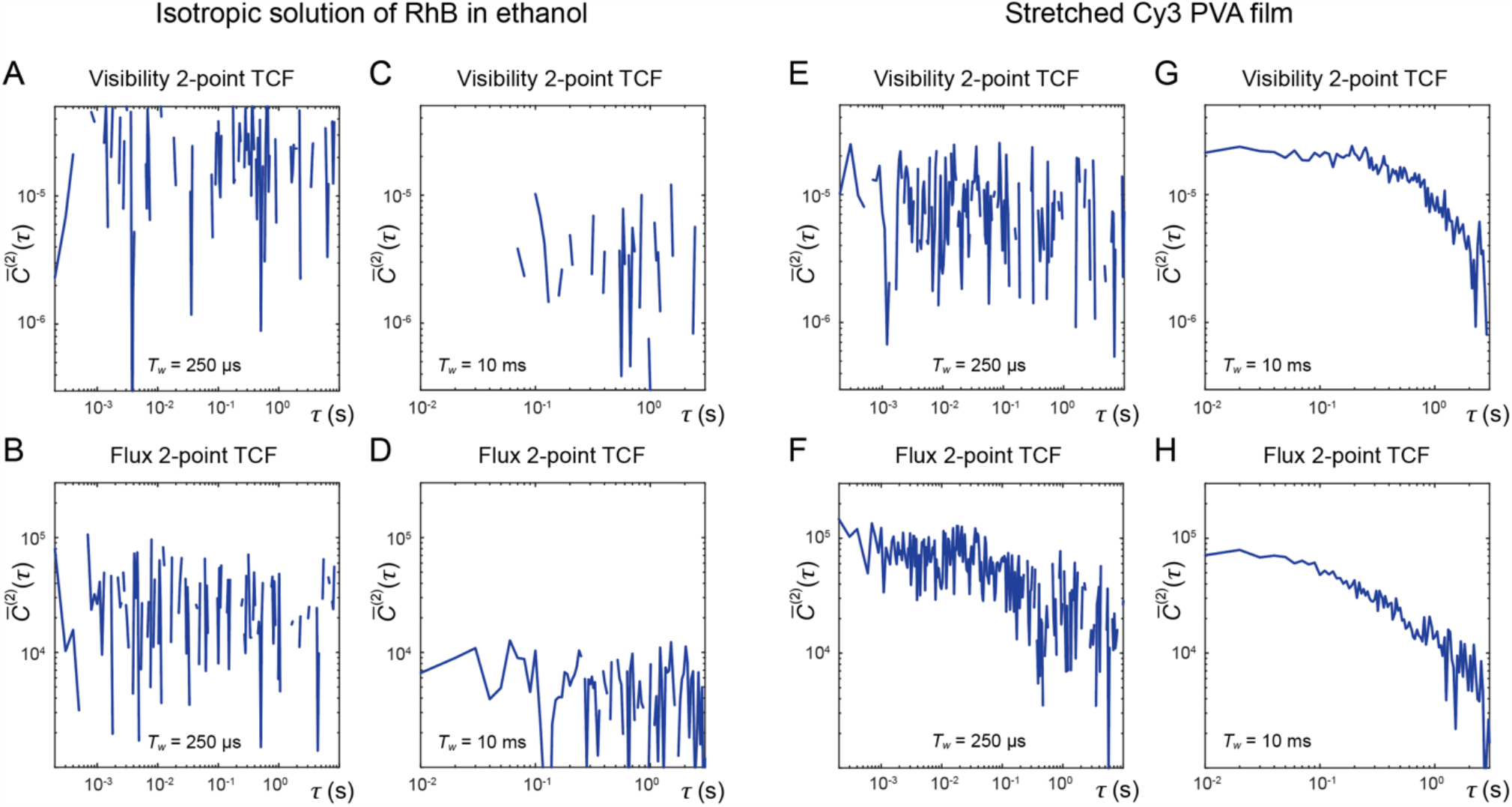
Control measurements of two-point time correlation functions (TCFs) for an isotropic solution of RhB in ethanol (panels ***A*** *–* ***D***) and for an anisotropic stretched Cy3 polyvinyl alcohol film (panels ***E*** *–* ***H***). Visibility and flux TCFs are shown in the top row (***A, C, E, G***) and the bottom row (***B, D, F, H***), respectively. Comparisons are shown using the two different integration windows *T*_w_ = 250 μs (***A, B, E, F***) and *T*_w_ = 10 ms (***C, D, G, H***). While the TCFs for the isotropic solution show no discernible decay, the TCFs for the stretched Cy3 PVA film exhibit a seconds-long decay component in both the visibility and flux.

